# Walking *Drosophila* navigate complex plumes using stochastic decisions biased by the timing of odor encounters

**DOI:** 10.1101/2020.03.23.004218

**Authors:** Mahmut Demir, Nirag Kadakia, Hope D. Anderson, Damon A. Clark, Thierry Emonet

## Abstract

Insects find food, mates, and egg-laying sites by tracking odor plumes swept by complex wind patterns. Previous studies have shown that moths and flies localize plumes by surging upwind at odor onset and turning cross- or downwind at odor offset. Less clear is how, once within the expanding cone of the odor plume, insects use their brief encounters with individual odor packets, whose location and timing are random, to progress towards the source. Experiments and theory have suggested that the timing of odor encounters might assist navigation, but connecting behaviors to individual encounters has been challenging. Here, we imaged complex odor plumes simultaneous with freely-walking flies, allowing us to quantify how behavior is shaped by individual odor encounters. Combining measurements, dynamical models, and statistical inference, we found that within the plume cone, individual encounters did not trigger reflexive surging, casting, or counterturning. Instead, flies turned stochastically with stereotyped saccades, whose direction was biased upwind by the timing of prior odor encounters, while the magnitude and rate of saccades remained constant. Odor encounters did not strongly affect walking speed. Instead, flies used encounter timing to modulate the rate of transitions between walks and stops. When stopped, flies initiated walks using information from multiple odor encounters, suggesting that integrating evidence without losing position was part of the strategy. These results indicate that once within the complex odor plume, where odor location and timing are unpredictable, animals navigate with biased random walks shaped by the entire sequence of encounters.

## INTRODUCTION

Olfactory search strategies depend on both an animal’s locomotive repertoire and the odor landscape it navigates. Navigational strategies have been investigated in a variety of odor plumes, each exhibiting a particular structure in space and time. The statistics of these plumes governs what information is available to the animal as it navigates, which in turn dictates the sequence of behaviors it can use to find its target. In some environments, such as the diffusion-dominated odor landscapes of *Drosophila* larvae, concentrations vary relatively smoothly from point to point. Accordingly, larvae can progress towards odor sources by sampling odor gradients spatially and temporally [2–5]. Similarly, adult flies in gradients can walk up the gradient by monitoring the odor intensity difference across their antennae pairs [6, 7].

In the absence of gradients, odor landscapes may still be relatively simple when the airflow is laminar. This is true for modest wind speeds and for homogeneous turbulent flows near smooth surfaces [8], and is often satisfied in laboratory wind tunnel experiments. In these conditions, odor concentrations might fluctuate, but on timescales that are not very short compared to the timescale of odor changes due to walking or flying. Furthermore, the largely unidirectional and steady wind provides a reliable cue about odor source location. In such environments, walking and flying moths accelerate and turn upwind upon entering the plume, and cast perpendicular to the wind or counterturn when the plume is lost [9–17]. Upwind turn responses and casting have also been observed in flying fruit flies as they navigate narrow odor ribbons [18, 19]. Similarly, flies walking in laminar flows turn upwind at the onset of spatially uniform plumes of natural [20, 21] or optogenetic fictive odors [22], and turn downwind or initiate a local search when the plume is lost. Likewise, in the laminar sublayer of turbulent flows, fluctuations in odor concentration can be relatively small, provided the source and average wind directions are not shifting and the surface is smooth [8]. Experiments in walking flies suggest that, in this case, high frequency fluctuations in odor concentration might be ignored and upwind progress may result from temporal integration of the odor concentration [20]. In all of these situations, either the spatiotemporal complexity of the odor signal is relatively low, odor encounters are relatively long, or both.

In contrast, odor landscapes are irregular in the bulk of turbulent flows [10, 23–26] or on rough surfaces where obstacles – like grass, shrubs and branches – and shifting winds introduce eddies and fluctuations that can perturb the boundary layer [27, 28]. Measurements of odor concentrations in forests [24, 29, 30] show that not only are local concentration gradients less indicative of odor source location, but importantly, odor encounters are intermittent, occurring as a random sequence of brief bursts. Theory suggests that in complex intermittent plumes, the timing of odor encounters may provide important information to the navigator [31, 32]. Indeed, moths follow tight trajectories upwind while navigating within a turbulent plume, much narrower than those in steady ribbons [9, 12, 15]. These narrow tracks were recapitulated for moths navigating pulsed ribbons, provided the pulse frequency was high enough, again implicating encounter timing in upwind progress [9].

There is an important distinction between experiments informed by steady odor ribbons versus those by spatiotemporally complex plumes. In steady ribbons, navigational behaviors can be tied to individual odor encounters because the location of the ribbon can be measured and the time-dependent odor signal perceived by the animal can be inferred from its trajectory. In spatiotemporally complex plumes, behaviors can at most be correlated with plume statistics, since the time when each individual filament hits the animal is unknown. In this context, analyzing how animals use the timing of individual encounters to navigate would require simultaneous measurement of behavior with odor.

Here, we investigated how the navigational strategies of freely-walking *Drosophila* are shaped by their encounters with individual odor packets in spatiotemporally complex odor plumes. We exploit a technical advance that relaxes the tradeoff between restricting odor dynamics and animal motion: an attractive odor that can be imaged in real time with unrestrained walking flies. By passing this odor in a laminar airflow and perturbing it with random lateral air jets, we generate a spatiotemporally complex plume whose statistics approximate those of turbulent plumes near boundaries [1]. This odor allows us to study walking fly olfactory navigation by directly connecting navigational behaviors to individual odor encounters.

Consistent with prior studies [33, 34], we find that flies on average walk upwind within the odor plume cone. However, upwind bias does not result from an accumulation of orientation changes following every odor encounter. Instead, flies execute stochastic, stereotyped 30-degree saccades at a rate independent of the duration or frequency of odor encounters. Upwind bias results not from modulating turn magnitude or frequency but rather turn direction: the randomly-occurring saccades are more likely to be oriented upwind when the frequency of odor encounters – but not their duration or concentration – is high, suggesting an important role for precise odor timing detection [35–38]. Prior studies have shown that flies increase walking speed at the onset of a uniform odor blocks [20, 39, 40]. In our spatiotemporal plume, flies spend only a fraction of time (~15%) experiencing detectable odor concentrations, and we expectedly do not find an appreciable increase in walking speed. However, flies do markedly modulate their rate of walking and stopping. In contrast to turn decisions, the rates of these walk-stop transitions are strongly tied to the frequency of encounters. We model stops and walks as a double, inhomogeneous Poisson process and find using maximum likelihood estimation and cross-validation that stop rates reset at every encounter before decaying back to a baseline rate. This suggests that individual encounters prolong the flies’ tendency to continue walking, but only for a brief time. Meanwhile, walks are triggered by accumulating evidence from multiple encounters while stopped. Using agent-based simulations, we show that this modulation of stops and walks shaped by the timing of odor encounters greatly enhances navigation performance. Together, our results suggest that navigation within spatiotemporally complex odor plumes is shaped by the sequence of encounters with individual odor packets. Both electrophysiological and behavioral measurements indicate that *Drosophila* – along with other insects, mammals, and crustaceans, among others – can precisely encode odor timing within their signal transduction cascade [35, 38, 41, 42]. Our findings suggest that *Drosophila* leverage this capability to navigate their olfactory world.

## RESULTS

### Visualizing dynamic odor plumes simultaneously with fly behavior

To investigate how freely-walking insects navigate odor plumes that are complex in both space and time, we developed a wind-tunnel walking assay for *Drosophila melanogaster* (Figure 1A). The large size of our 2D arena (300 x 180 x 1 cm) allowed us to simultaneously image several flies in the dark with minimal mutual interactions. The main flow was set to 150 mm/s, chosen as sufficiently strong for flies to tax upwind, but not so strong that they remained stationary [43]. Plumes that fluctuated in space and time were generated by injecting odors at the center of an air comb and perturbing the laminar flow with lateral jets stochastically alternating at a Poisson rate of 10/sec. To visualize the flow, we injected smoke, which is turbid, into the center of the air comb and imaged it in the infrared at 90 Hz. Serendipitously, we noticed that when we placed starved flies in the assay with the fluctuating smoke, flies walked upwind toward the source (Figure 1B, Video S1) in a manner reminiscent of their behavior when we injected an attractive odor such as ethyl acetate. We reasoned that if this attraction to smoke were olfactory, the imaged smoke intensity could then provide a proxy for odor concentration, allowing us to visualize dynamic odor plumes simultaneously with fly behavior (Figures 1C-E and Video S2).

**Figure 1:**
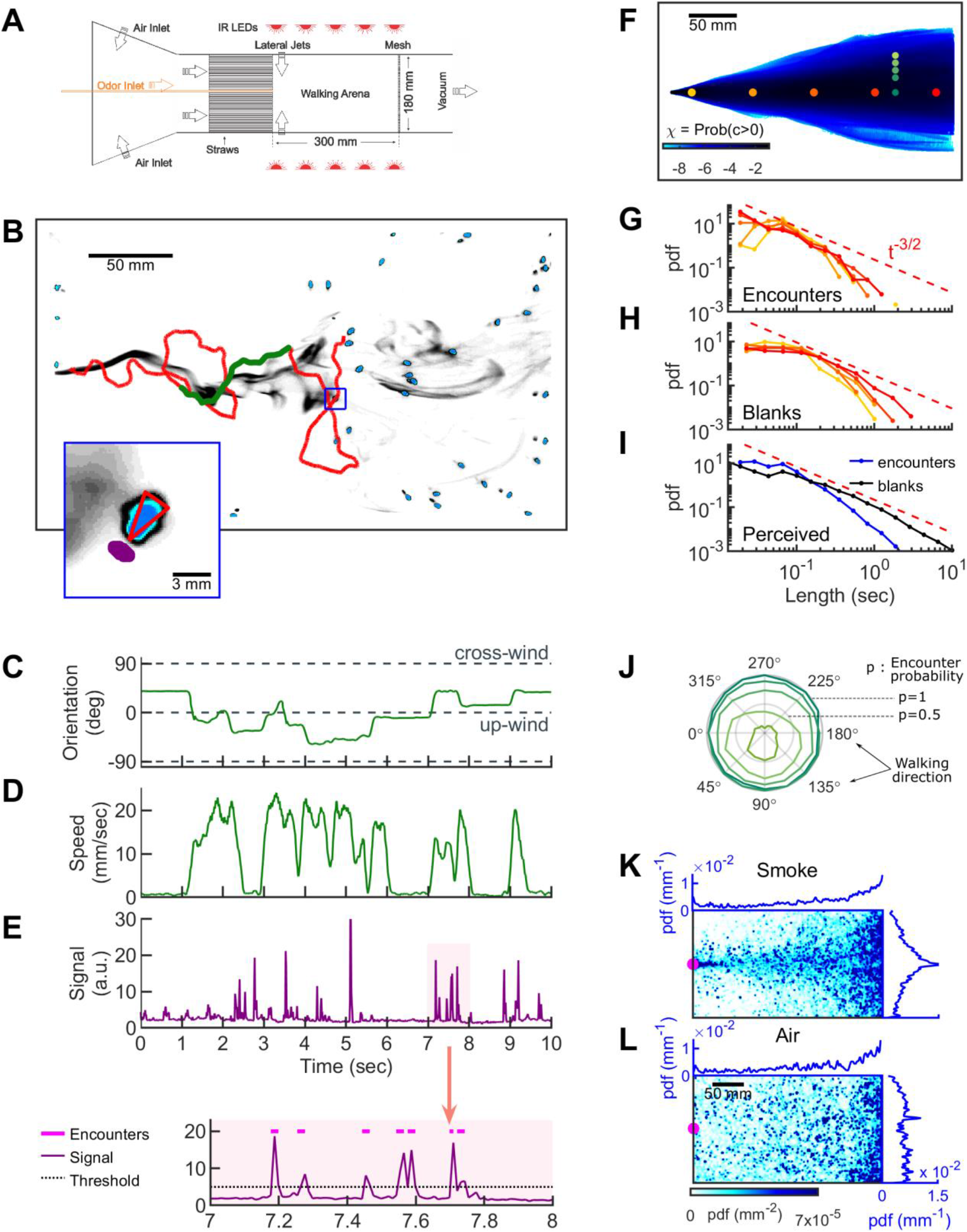
Simultaneous visualization of odor and fly behavior. **A**. Experimental apparatus. The main flow (150 mm/s) is perturbed by lateral jets (1500 mm/s) that alternate stochastically with characteristic time scale 100 ms. **B.** Walking arena with flies (blue), odor intensity (grey), and a representative trajectory of a navigating fly (red). Green: portion of the trajectory plotted in C-E. Note that since the odor environment is fluctuating, the image in (B) only represents the environment at a random time point. Blue rectangle: area shown in the inset. Inset: blue: fly; grey: odor intensity; red triangle: orientation of the fly; purple: the virtual antenna in which the odor intensity is averaged as a proxy for the signal perceived by the fly. Simultaneous orientation (C), speed (D), and perceived stimulus (E) of the fly while it is navigating in the intermittent plume (green portion of the trajectory in (B)). Orientation and speed were smoothed with a 100 ms sliding box filter. Shaded area in E (top) is plotted at a larger scale (bottom) with the threshold (dotted line) used to identify the odor encounters (magenta lines above the signal trace in purple) (see also Figure S2 and Methods). **F.** The fraction of time that odor concentration is above zero (i.e. intermittency) at fixed locations. Image intensity is median-filtered (square filter size 2.3 mm), and the likelihood that the intensity is above zero, averaged over all frames of the video. Red dots: positions of encounter and blank duration distributions plotted in G-H. Green dots: positions of encounter probability distributions plotted in J. **G-H** Distributions of encounter and blank durations, respectively, at positions color coded in F. The red dash line shows the *t*^−3/2^ expected from theory for a turbulent flow [1]. **I.** Odor encounter and blank durations experienced by navigating flies **J.** Probability (r axis) to have an odor encounter within 1 sec while walking with a speed of 10 mm/s, starting from positions color coded in F, as a function of walking direction (theta axis). **K.** Probability distribution functions (pdf) of fly positions in the arena for the complex smoke plume as in A (n=1073 trajectories). **L.** Same without smoke but with the same complex wind pattern as in K (n=502). Magenta: location of the source. Blue curves: marginal pdfs.

Smoke is a complex stimulus (Figure S1A), containing not only CO_2_ and volatile chemicals, but also heat, humidity, and airborne particles. We therefore set out to verify that the attraction to smoke is olfactory. For this purpose, we used a simplified environment consisting of a standing odor ribbon, which we generated in our assay by maintaining the laminar flow and odor injection but turning off the lateral jets (Figure S1B). First, we compared behavioral statistics in smoke to those in the attractive odors ethyl acetate (EA) and apple cider vinegar (ACV). The likelihood that flies were in the narrow band near the smoke ribbon increased with smoke concentration (Figure S1C), before saturating at a sufficient dose (Figure S1D), a result reproduced in both EA and ACV (Figures S1E-H). We then tested contributions from carbon dioxide sensing and vision, using Gr63a^-/-^ [44] and norpA^-/-^ [45] mutants, respectively. Both mutants retained the ability to localize the odor source at a level comparable to wild-type flies (Figure S1IJ). To test whether humidity played a major role we saturated the airflow with 80% humidity and found that source localization was reduced but still significantly above random (Figure S1IJ). Finally, we tested the olfaction directly using Orco^-/-^ mutants [46], as well as anosmic flies (Gr63a^-/-^, Orco^-/-^, Ir8a^-/-^, and Ir25^-/-^) [47]. In both sets of mutants, the ability to find the odor source was completely abolished. Orco^-/-^ mutants (but not the anosmic flies) exhibited a slight repulsion to the smoke ribbon, which we attributed to an aversive response to carbon dioxide (Figure S1IJ) [46, 48]. Thus, flies’ attraction to smoke is driven mainly by olfaction.

To quantify the time-dependent stimulus experienced by each fly during navigation we averaged the signal intensity in a small area (1.10 mm^2^) near its antennae (Figures 1B inset, 1E, and Video S2). The onset and offset of odor encounters were defined as the times when the signal crossed a threshold, which we set to 2.5 SD (σ) above the background noise. We refer to the periods when the odor is above threshold as “odor encounters,” or “encounters” for short, and periods when the odor is below threshold as “blanks”. We verified that the results and conclusions presented below remained unchanged for thresholds between 2.0σ and 3.5σ. Using the 2.5σ threshold, the error in the timing of odor encounters was estimated to be less than 25ms (Figure S2 and Methods). Using this setup, we then examined how walking flies navigate odor plumes that fluctuate in both space and time.

### Within the odor plume, odor encounters are brief, frequent, and unpredictable

We first quantified the statistics of the odor environment the flies must navigate. Our odor plume was highly intermittent, composed of spatiotemporally localized filaments breaking continuously in time (Figure 1B and Video 1, lateral jets on). Odor intermittency – the fraction of time the odor was above sensory threshold – ranged several orders of magnitude in the conical extent of the plume (Figure 1F). In general, intermittency was very low and reached 19% only very close to the source. At fixed locations from the source, odor encounter (Figure 1G) and blank (Figure 1H) durations spanned a wide range of time scales, approximately approaching the power law ~ *t*^−3/2^ theoretically predicted for turbulent odor plumes in the atmospheric boundary layer. The distribution of encounter and blank durations experienced by navigating flies spanned an even greater range and were closer to a power law (Figure 1I). On average, flies experienced brief odor encounters (mean duration ~200ms) at a mean frequency of 4Hz. Even beyond their variability and brevity, encounters were also highly unpredictable in location. To quantify this, we calculated the likelihood to receive an odor encounter in 1 second, assuming one walks straight at 10 mm/s radially outward from a fixed point. Predictability in the location of future odor encounters would then manifest as a directional dependence of this likelihood. Within the conical extent of the plume, the likelihood was nearly isotropic with respect to walking direction, whereas near the plume edges, likelihoods were skewed towards centerline of the plume cone (Figure 1J). Within the conical extent of the odor plume, therefore, the location of future odor encounters was uncertain. Despite this uncertainty, flies remained largely in the plume cone and were able to successfully locate the odor source, whereas when in the fluctuating wind only, they could not locate the source (Figures 1K-L).

### Stopping and turning comprise the bulk of the navigational repertoire within a spatiotemporally complex odor plume

How are fly orientation and speed shaped on average by an odor signal exhibiting this degree of spatiotemporal complexity? To compare these behaviors to those in an odorless environment, we presented the complex plume in 15-second blocks by closing and opening the odor valve every 15 seconds, but maintaining the alternating lateral jets throughout the trial. This produced an environment in which a 15-sec block of complex odor plume alternated with a 15-sec block of fluctuating wind only (Videos S3). When the odor was on, odor encounters were frequent, but randomly experienced in time (Figures 2A-B). As expected, flies were more likely to be oriented upwind when the odor was on (Figure 2C), as previously reported [14, 18, 20–23, 49]. However, unlike for flies walking into a spatially homogeneous odor block [20, 39, 40], changes in average angular speed were minor, with a less than 10% change between blocks (Figure 2D). Walking speeds were similarly unmodulated, again in contrast to walking flies in homogenous odor blocks (Figure 2E) [20]. This is not inconsistent, however: since encounters were so brief (~200 ms), the integration timescales for speed modulation measured previously [20] would only produce <10% increase in either ground or angular speed.

**Figure 2:**
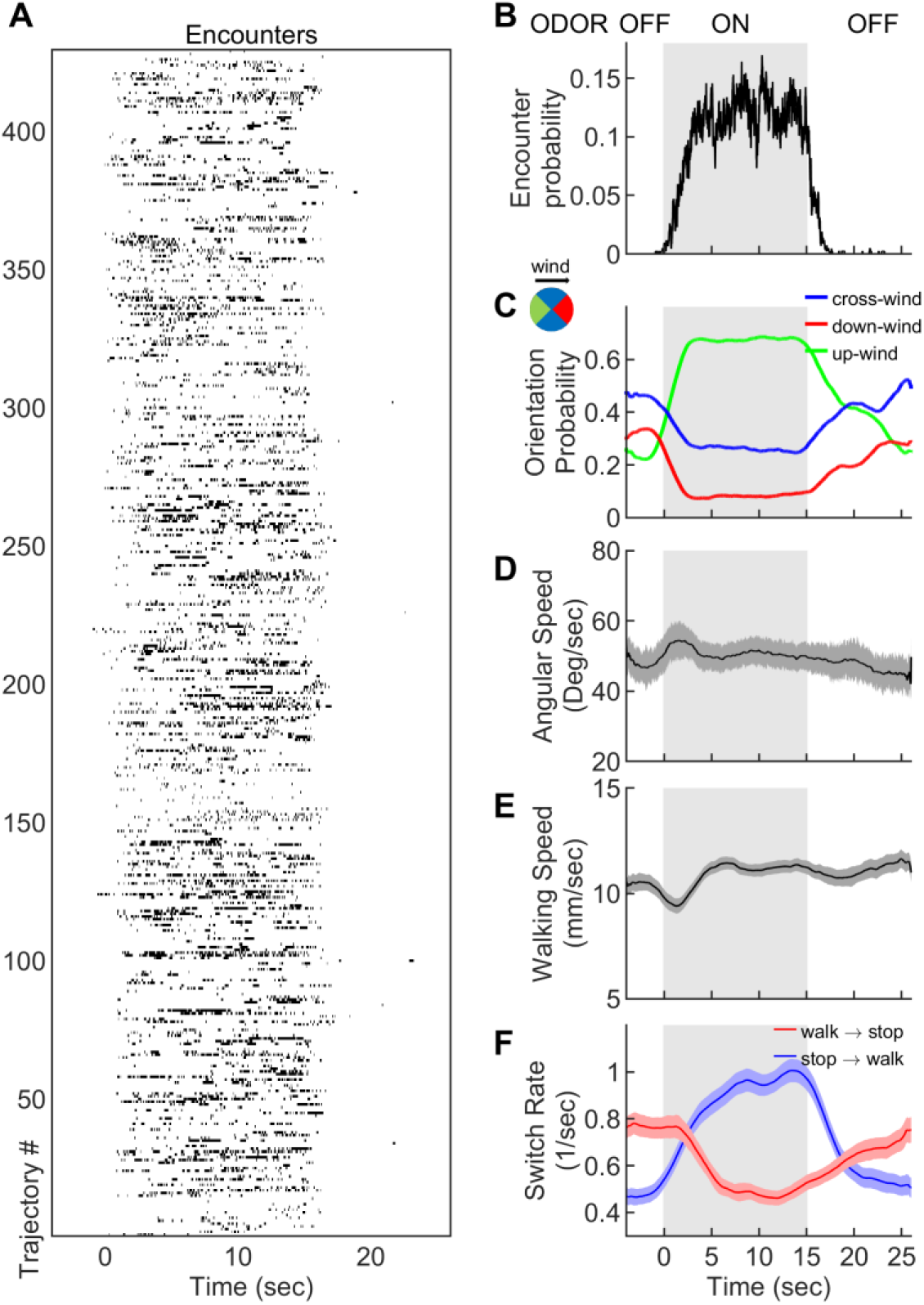
Navigation within intermittent complex odor plumes comprises walk-stop transitions and upwind orientation. **A.** The odor encounters experienced by flies navigating the intermittent smoke plume as in Figure 1B. While the wind is continuously perturbed by the lateral jets, the odor is turned on/off in 15 sec blocks (grey shading). Rows indicate independent trajectories (*n* = 429) obtained from 267 flies. **B-F.** Quantities averaged over all trajectories in A as a function of time. **B.** Probability of having an encounter. **C.** Probability of being in up-wind (green), cross-wind (blue) and down-wind (red) orientations estimated in 90 degree quadrants as shown in the circle with the same color codes. C-E include only time points for which the fly was walking (*v* > 2 mm/s). **D.** Angular speed. **E.** Walking speed. **F.** Walk-to-stop (red) and stop-to-walk (blue) switching rates. All quantities in B-F are smoothed with a 5-second sliding box filter. Error bars indicate SEM.

Though changes in ground speed were minor, we noticed a high incidence of stopping in our spatiotemporally complex plume (Figure 1D). The prevalence of immobility has been noted before in walking flies navigating homogenous odor blocks [20], though its role in navigation was not investigated. We suspected that stopping might form a critical component of intermittent plume navigation for walking flies. Indeed, walk-to-stop and stop-to-walk transition rates were strongly modulated during the transitions between odorized and non-odorized blocks (Figure 2F). Natural odors ACV and EA elicited similar navigational trends in angular and ground speeds, orientation, and stopping rate when presented in these 15-second blocks (Figure S3). Together, this suggested that turning and stopping comprised the bulk of the navigational repertoire for walking flies in spatiotemporally complex plumes. This prompted us to next examine how the sequence of individual odor encounters experienced by navigating flies precisely shapes their decisions to turn, walk, and stop.

### Upwind orientation results from repeated odor encounters, not cumulative odor exposure time

Flies reorient upwind soon after flying into an odor ribbon [19] or walking into a homogeneous odor block [20]. For this reason, we calculated the change in fly orientation following an individual encounter. We found that within two seconds of an encounter onset, flies of any orientation biased their orientation upwind (Figures 3A and S4A–S4B). Since encounter frequency was on the order of a few Hz, flies receiving one encounter were likely to receive more within the 2 second window. Upwind bias may therefore reflect an accumulated effect from repeated odor encounters. Partitioning the data into encounters followed by 0, 1-3, or 4+ further encounters within two seconds, we found that odor encounters followed closely by many others elicited much stronger upwind bias than did isolated ones (Figure 3B). To quantify this more precisely, we calculated a running average of encounter frequency *W_freq_*(*t*) by convolving the binary vector of encounter onset times with an exponential filter (timescale *τ* = 2s), and plotted upwind orientation as a function of encounter frequency (Figure 3C). All orientations were reflected over the *x*-axis, whereby 0° is upwind and 180° is downwind. The trend was strongly monotonic, with an intercept of 88.6° at 0 Hz – flies experiencing no encounters were oriented nearly equally upwind and downwind – and a slope of 21.6°/Hz (*p* < 10^-4^) – flies experiencing a frequency of 3 Hz would be oriented just 25° off the upwind direction. If no further encounters were received, this monotonic trend dropped steadily to a slope of 4.5°/Hz (not significantly different from 0, *p* > 0.05) after 5 seconds (Figure 3C). This suggests that repeated interactions with the plume biased the fly upwind, and after some time without encounters, flies were again uniformly oriented.

**Figure 3:**
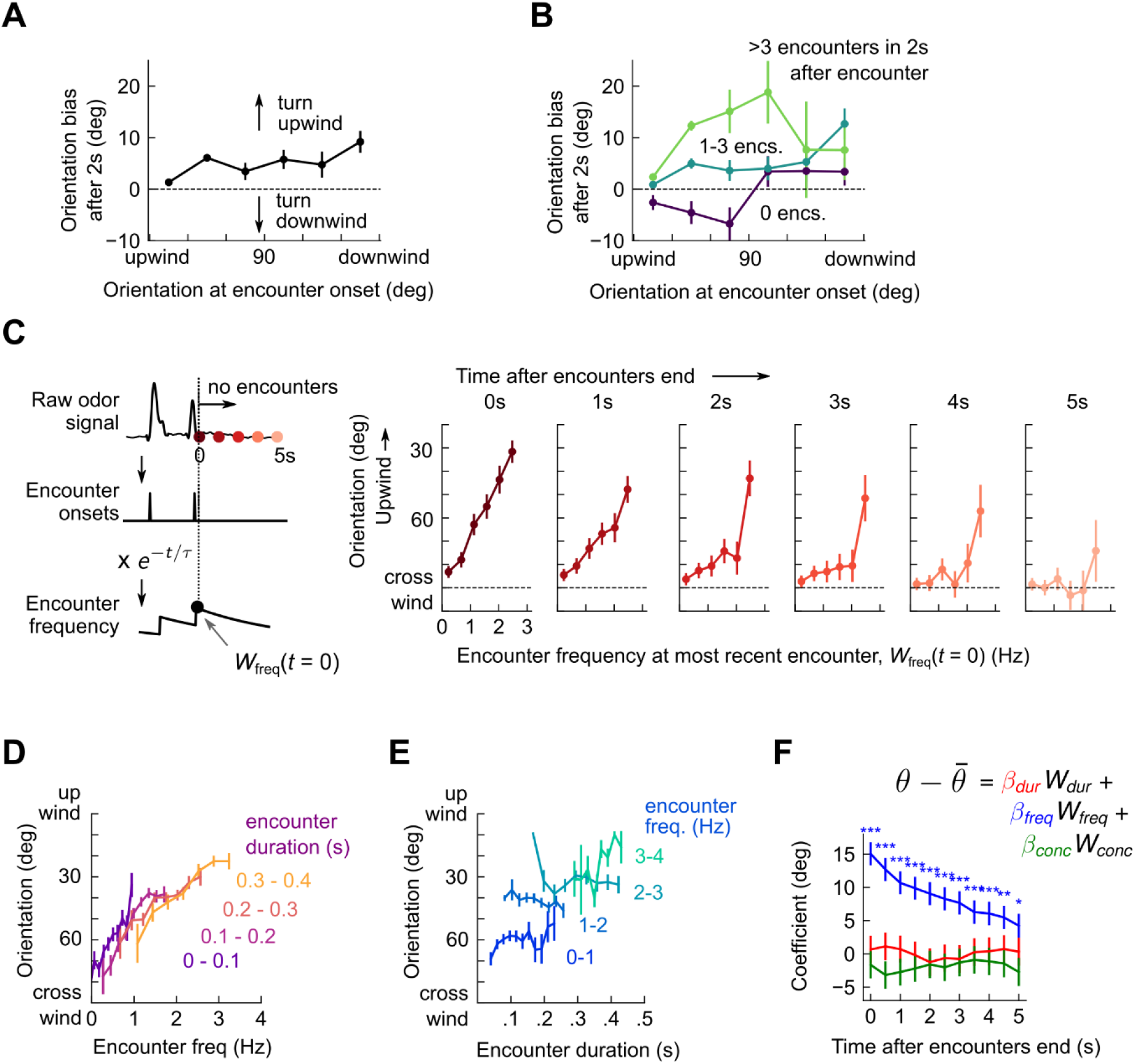
Flies use encounter frequency to bias orientation upwind. **A.** Orientation 2 seconds after an encounter, as a function of orientation at encounter onset (*n*=5040). The mean orientation change for random times is subtracted out. (right) **B**. Same data, now binned by number of subsequent encounters in the 2s window. **C.** Orientation as a function of encounter frequency at the most recent encounter, *W_freq_*(*t* = 0), for various times *t* > 0 after encounters have been interrupted. Encounter frequency is defined by convolving the binary vector of encounter onsets with an exponential filter of timescale *τ* = 2s. *t* = 0 defines the time of the most recent encounter, and the individual plots show fly orientation as a function of the encounter frequency at this time, *W_freq_*(*t* = 0). Orientation biases strongly upwind with frequency, and this correlation vanishes after ~5 seconds. **D**. Orientation versus encounter frequency when encounter duration is held fixed within a small range, for various ranges. **E**. Orientation versus encounter duration when encounter frequency held fixed within a small range. **F**. Estimated regression coefficients for a trilinear fit of fly orientation to encounter frequency, encounter duration, and signal intensity. Each of the independent variables has been standardized. Coefficients are plotted for various times after encounters are interrupted (as in C). Statistical significances using a 2-tailed t-test are shown next to curves; if no stars are shown, the coefficients are not statistically distinct from 0. The data indicate that orientation correlated with encounter frequency, but not encounter duration or signal intensity.

The amount of time a fly is exposed to odor increases with each subsequent encounter. Does upwind bias result from the number of individual odor interactions, the cumulative duration of these encounters, or both? If, for example, all encounters were 200 ms long, then tripling encounter frequency would also triple perceived odor duration – frequency and duration would be perfectly correlated. But if orientation depended on odor duration alone, the dependency on frequency noted above would arise simply as a consequence of this correlation. Prior results suggest that walking flies bias orientation and speed by filtering odor in time [20], so we suspected that odor duration might contribute to some or all of the upwind bias. To investigate this possibility, we defined a running average of odor duration *W_dur_*(*t*) analogously to *W_freq_*(*t*), by exponentially filtering the binary vector of odor intermittency (1 during encounters, 0 during blanks). We disassociated Wfreq(t) and *W_dur_*(*t*) by holding one constant to a small range, and plotting upwind orientation against the other. Surprisingly, with this analysis, only the correlation of orientation with encounter frequency remained (Figure 3D–3E). We also investigated the possibility that odor concentration contributed to upwind turning by defining *W_conc_*(*t*) analogously using the raw signal. While we have not quantified the exact relationship between odor concentration and image intensity, our dose response results (Figure S1C–S1H) suggest that they are monotonically related, so a correlation would exist to first order. Linearly regressing upwind orientation simultaneously against *W_freq_*(*t*), *W_dur_*(*t*), and *W_conc_*(*t*), revealed *W_freq_*(*t*) as the sole explanatory variable (*p* < 1e-6, *p* > 0.05, *p* > 0.05, respectively) (Figure 3F). Together, these results indicate that in the intermittent, spatiotemporally complex plumes in this experiment, upwind orientation was driven by the frequency, but not by the duration or concentration, of odor encounters.

### Odor encounters bias turn direction, but not turn likelihood or turn magnitude

The lack of a clear upwind bias following an isolated encounter (Figures 3B and S4B) suggested that reorientations may not be simply an encounter-elicited reflex. To characterize reorientation, we first thresholded angular speed to identify turn events (Figure S5, and methods). We found that individual turns occurred not in a continuum of angles but rather in discrete saccades of 30°±10° either left or right (Figure 4A), consistent with previous studies in unodorized environments [50]. Moreover, the contribution to upwind bias from the inter-saccade sections of the trajectories was not significant (Figure 4B). This indicates that the discrete saccadic turns were responsible for upwind progress during navigation. The waiting time between saccades obeyed an exponential distribution with timescale *τ* = 0.75s±0.17, or a Poisson rate of about 1.3 turns per second (Figure 4C). Surprisingly, this turn rate was insensitive to either encounter frequency or duration (Figure 4C). This presented a puzzle: if flies turned left and right at discrete angles and a constant rate, they were effectively executing a random walk on the circle. Since angular random walks randomize orientations in time, how would flies orient upwind? Partitioning the turn angle distribution into bouts of low (< 1 Hz) and high (> 4 Hz) encounter frequency resolved this puzzle. For high frequencies, the distribution of turn angles exhibited the same ±~30° peaks, but now with an upwind lobe much larger than the downwind one (Figure 4D). Thus, odor encounter frequency biased the direction of turns, while leaving the magnitude and rate of turns unchanged.

**Figure 4:**
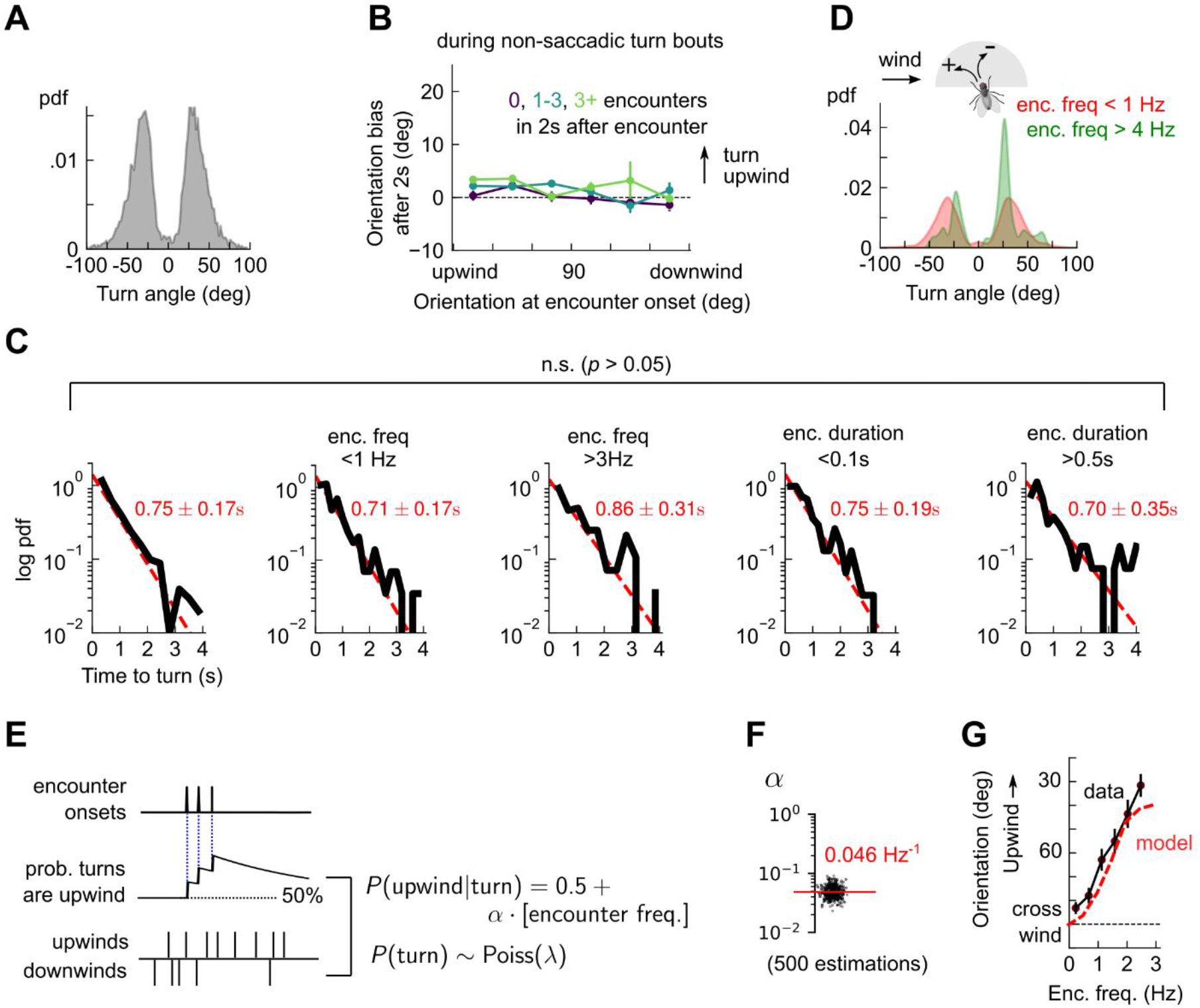
Odor encounters bias turn direction, but not turn rate or turn magnitude. **A.** Distribution of change in orientation following a turn. The discreteness of turn angle (two narrow peaks in pdf) was verified to be insensitive to the threshold used to determine turns (Figure S5). **B**. Cumulative change in orientation over non-turning bouts (“straight” segments) 2 seconds after an encounter, versus orientation at encounter onset (compare with Figure 3B). Data are partitioned into encounters followed by 0, 1-3, or 4+ subsequent encounters in the following 2 seconds. **C**. Leftmost plot: Distribution of times until a turn. Red line: maximum likelihood fit to exponential distribution, with mean 0.75 ± 0.17s (distribution generated by bootstrapping). Remaining plots: Same, now for times at which encounter frequency is low (<1Hz; 2^nd^ plot) or high (>3Hz; 3^rd^ plot), or times at which encounter duration is low (<100 ms; 4^th^ plot) or high (>500 ms; 5^th^ plot). Fits are 0.71 ± 0.17s, 0.86 ± 0.31s, 0.75 ± 0.19s, 0.70 ± 0.35s, respectively, none of which are statistically distinct from the data for all turns (*p* > 0.05, 2-tailed t-test). **D**. Distribution of turn angles during low (< 1 Hz) or high (> 4 Hz) encounter frequency bouts (compare A). **E**. Model of fly turning. Turn events obey a Poisson process with timescale *τ_T_* = 0.75s (as in C). Turn direction is chosen randomly at each turn time; the probability *p_T_* that the turn is directed upwind increases at every encounter by a factor *α*, before decaying back to a baseline of 0.5, modeled by 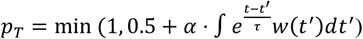, where *τ* = 2s (Methods). **F**. Distribution of gain parameter *α*, estimated for 500 distinct subsets of the data. The distribution is highly peaked, indicating its robustness. **G**. Upwind orientation versus encounter frequency for data (black) and model prediction (red).

These findings could be recapitulated with a simple stochastic model of turning, in which walking flies execute Poisson turns at a constant rate. The magnitude of each turn is chosen randomly from the measured distribution, and the likelihood *p* that the turn is directed upwind increases by a gain factor of *a* at every encounter onset, before decaying to a baseline of *p* = 50% (Figure 4E). We estimated the parameters from a maximum likelihood fit to the data, obtaining a distribution of parameters by performing the estimation on 500 distinct subsets of the measured data. The distribution of estimated gains *a* clustered tightly around a mean of *a* = 0.046 1/Hz (Figure 4F), indicating that the parameter estimates were robust. Simulating this model with the mean of the estimated parameters closely reproduced the dependence of upwind orientation on encounter frequency (Figure 4G). Together, these findings indicated that in the spatiotemporally complex plume, odor encounters did not initiate reflexive upwind turning. Rather, odor encounters increased the likelihood that stochastically-occurring, saccadic left/right turns were directed upwind.

### Stopping and walking are stochastic events whose rates depend on odor encounter timing

Walking flies navigating spatiotemporally complex plumes stopped frequently (Figure 1D), and the rate of both stopping and starting depended strongly on the presence of odor (Figure 2F). To connect walk-stop transitions to individual encounters, we first calculated the likelihood to be walking or stopped during the 2 seconds after an encounter (Figure S6A). Walking flies were more likely to remain walking after an encounter (versus random times), while stopped flies were more likely to initiate a walk. Notably, even a single encounter was sufficient to initiate walks, and higher encounter frequencies biased this further (Figure S6B). This implicated both individual encounters and encounter history in decisions to walk or stop. In contrast, we found no change in walking speed following encounters (Figure S6C), even when encounter frequencies were appreciable (Figure S6D).

How does the sequence of encounters shape a fly’s decision to walk or stop? After an odor encounter, flies walked for longer periods before stopping, compared to random (Figure 5A). Thus, encounters reduce stopping likelihood, and flies experiencing higher encounter frequencies walked for longer (Figures 5B–5C). In addition, the time to stop following an encounter was the same, whether the encounter was isolated or part of a clump containing 3+ encounters in 1 second (Figure 5A). The times to stop were approximately exponentially distributed. We therefore modeled stop decisions as a Poisson process with time-dependent stopping rate *λ_w→s_*(*w*(*t*)), where *w*(*t*) is the binary vector of encounter onset times. We considered various models for the dependency of the stopping rate on the encounter sequence *w*(*t*). In the *last encounter* model, *λ_w→s_*(*t*) drops to the same given value at each encounter, before decaying back to baseline with some characteristic time *τ_s_* (Figure 5D). In the *accumulated evidence* model, *λ_w→s_*(*t*) decreases further at every odor encounter, and therefore remains at a lower value when encounters are more closely spaced (Figure 5H). In the *encounter duration* model, *λ_w→s_*(*t*) switches between a low value during encounters and a higher value during blanks (Figure 5I). These models contain various parameters dictating the baseline rates and timescales, which we fit to the data using maximum likelihood estimation. As in the turn model, we obtained a distribution of parameters by carrying out the estimation on 500 distinct subsets of the data – quantifying the robustness of each parameter (Figure S7A).

**Figure 5:**
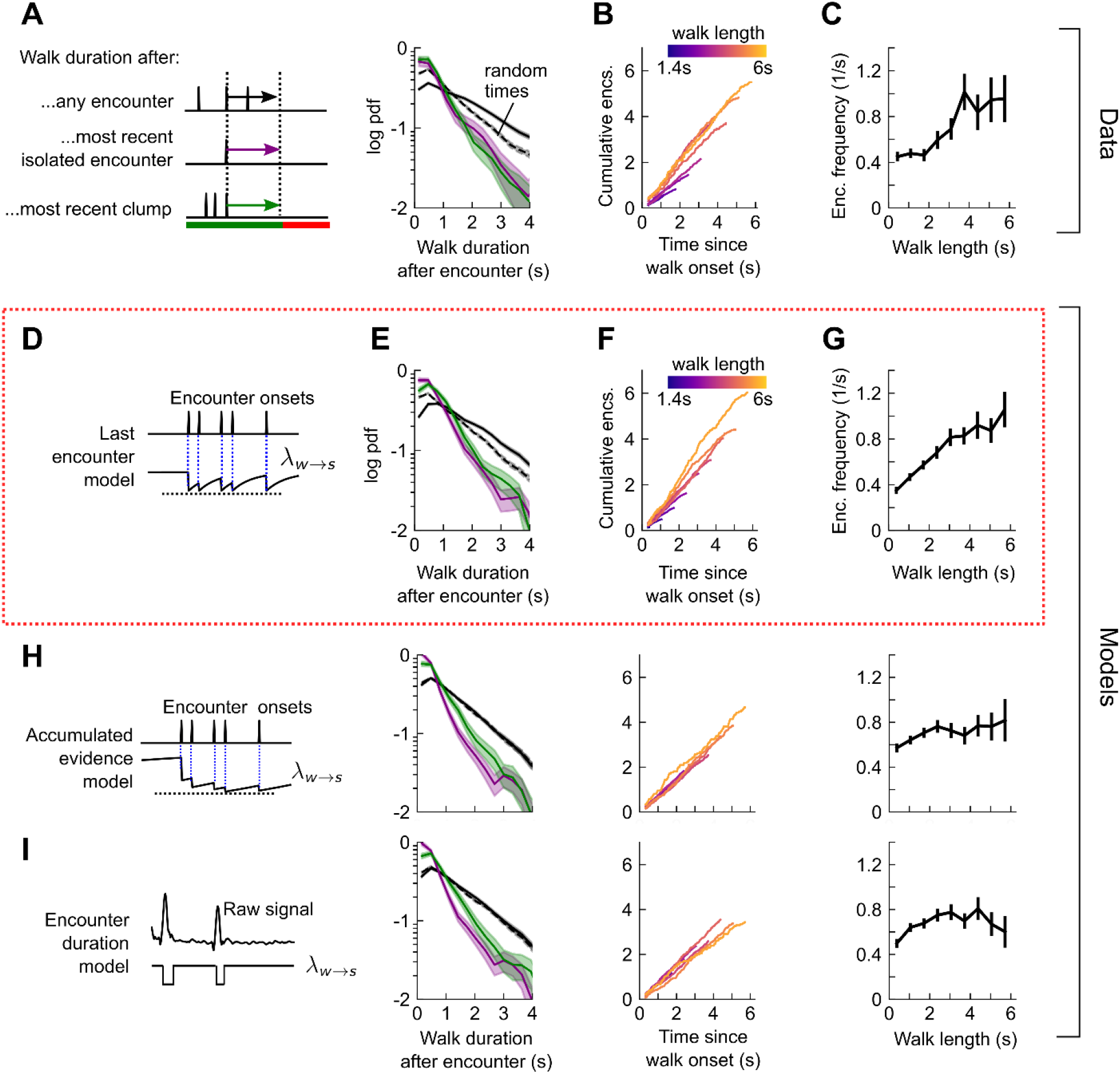
Stops decisions are stochastic events whose rate is modulated by the timing of the most recent encounter. **A.** Distribution of walking duration following any odor encounter, or the most recent encounter or clump before a stop. Dotted line: distribution for randomly chosen times. **B.** Cumulative encounter counts since walk onset, for various walk durations. **C.** Average encounter frequency versus duration of walk bout. **D.** In the last encounter model, stop decisions are modeled as a Poisson process with inhomogeneous rate 2_*w→s*_(*t*), where 2_*w→s*_(*t*) resets to a fixed value at every encounter, then decays back to baseline. This is modeled by 2_*w→s*_(*t*) = *λ*_0_ + Δ*λe*^-Δ*T*(*w*(*t*))/*τ_s_*^, where Δ*T*(*w*(*t*)) is the time since the most recent encounter (Methods for details). Median of estimated parameters are λ_0_ = 0.78s^-1^, Δ! = −0.61s^-1^, *τ_s_* = 0.25s (Figure S7). **E-G.** Analogues of A-D using data generated by the model. **H.** Analogues of E-G for the accumulated evidence model. In this model, λ_*w→s*_(*t*) decreases at every encounter, but remains at a lower value when encounters are more closely spaced. We model this with 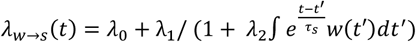. **I**. Analogues of E-G for the encounter duration model, in which 2_*w→s*_(*t*) switches between a low value during encounters and a higher value during blanks.

We found that the time-to-stop statistics were explained well by the last encounter model (Figure 5E-G), but not by the other models (Figure 5H-I). Our parameter fits indicate that at each encounter, the stopping rate drops to 0.17 s^-1^, before rising with timescale 0.25s to a background rate of 0.78 s^-1^. In the accumulated evidence model, the distribution of parameter estimates were broad, and often the parameters were estimated close to the imposed bounds (which ranged 2 orders of magnitude), suggesting that this model was not robust to the data (Figure S7B). The parameters of the encounter duration model were narrowly peaked, but the predictions were poor, so the model was incomplete (Figure S7C). In both these latter models, the distributions of walk durations following encounters was not higher than those following random times (compare black and dotted black lines in Figure 5A and 5E to those in Figures 5H–5I). Thus, our data indicate that flies continuously adjust their likelihood to stop while navigating, and that the rate of stops decreases by a factor of nearly 5 at the onset of each encounter. This decrease in stop rate at each encounter is brief, less than 1 second, suggesting that the ongoing perception of frequent encounters retains the flies in an active, walking state, but when encounters are interrupted, stops are frequent.

We next quantified the rate of stop-to-walk transitions. In contrast to stops, the time to walk was significantly shorter following a clump of encounters than an isolated encounter (Figure 6A), implicating the history of encounters in walk initiation. In addition, the cumulative number of encounters received during a stop bout was rather independent of stop length, ~ 0.75-1.25 encounters for stops between 2 and 6 seconds long (Figure 6B). This observation would not rule out models in which the walking likelihood accumulated with every encounter, nor those in which the rate jumped to a large, fixed value at each encounter. Therefore, we modeled walk decisions with 3 models analogous to those used for the stop decisions. In the accumulated evidence model, the rate increases by the same amount at each encounter (Figure 6D), while in the last encounter model, the walk rate *λ_s→w_*(*t*) increases to a set value at each encounter, before decaying to baseline (Figure 6H). In the accumulated evidence model, stopped flies receiving a clump would initiate walks sooner than those receiving a single encounter. In the encounter duration model, the rate switches between a higher value in encounters and a lower value in blanks (Figure 6I). The time-to-walk statistics were fit well by the accumulated evidence model (Figure 6E-G), but not the other two models (Figure 6H–6I). The estimated baseline walking rate is *λ*_0_ = 0.29 s^-1^, so stopped flies will on average remain stopped for ~3 seconds if they receive no signal. This rate increases at each encounter by Δ*λ* = 0.41s^-1^, before decaying to baseline *λ*_0_ with timescale of 0.52s. Though our model predicts that a higher frequency of encounters will elicit an earlier walk, Δ*λ* is comparable to the base rate – more than doubling the transition rate – so even a single encounter is sufficient to elicit a walk, as observed in Figure 6B). Together, this suggests that stopping forms a key component of the navigational strategy, and that stop and walk decisions are stochastic events whose rates of occurrence depend on the precise timing of recent encounters.

**Figure 6.**
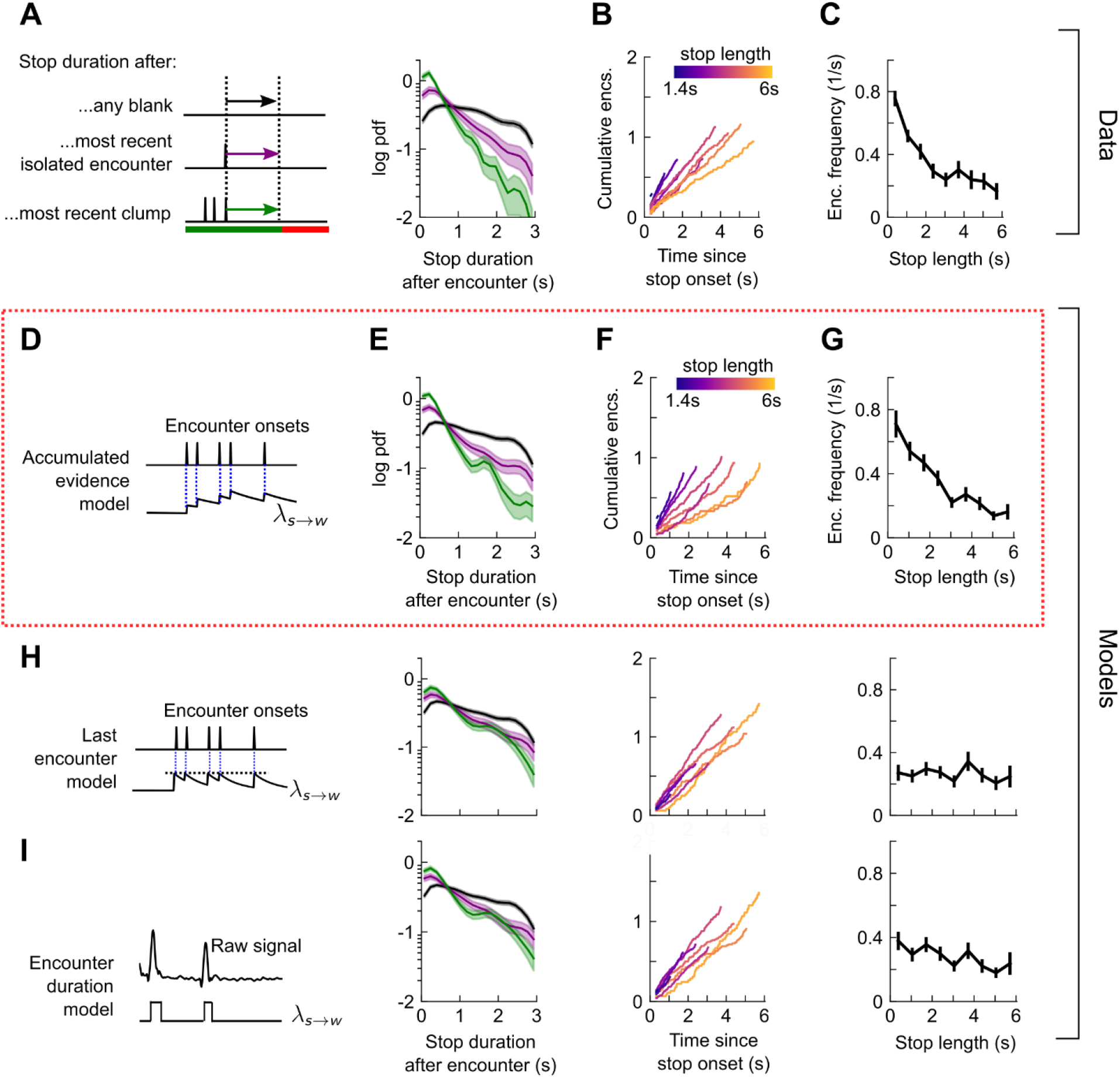
Walk decisions are stochastic events whose rate accumulates evidence from recent encounters. **A.** Distribution of walking duration following any encounter, or the most recent encounter or clump before a walk. **B.** Cumulative encounter counts since walk onset, for various stop durations. **C.** Average encounter frequency versus duration of stop bout. **D**. In the accumulated evidence model of walk decisions, *λ_s→w_*(*t*) increases at every encounter onset, before decaying to baseline. This is modeled by 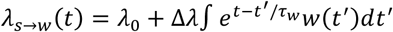. Median of estimated parameters are *λ*_0_ = 0.29s^-1^, Δλ = 0.41s^-1^, *τ_w_* = 0.52s (Figure S7). **E-G**. Analogues of A-C using data generated by the model. **H**. Analogues of E-G for the last encounter model, in which the walk rate increases to a fixed value at each encounter before decaying to baseline. **I**. Analogues of E-G for the encounter duration model, in which *λ_s→w_* (*t*) switches between a high value during encounters and l ow value during blanks.

### Encounter-modulated decisions enhance navigational performance

To test how these behavioral algorithms individually affected navigational performance, we incorporated our findings into an agent-based simulation (Figure 7A). We simulated 10,000 individual virtual flies navigating using the turn, stop-to-walk, and walk-to-stop models that we found in our data. In these simulations, we calculated both the likelihood that agents reach the source as well as the time taken to do so. Virtual flies implementing all three encounter-modulated behaviors navigated largely in the plume cone (Figure 7B) and converged to the source (Figure 7C), similarly to real flies (10.5% of real flies and 7.8% of virtual agents reached within 15 mm of the source). Visually, the simulated tracks resembled the measured tracks, containing non-linear, circuitous routes toward the source, as well as wide loops (Figure 7B). To meaningfully test the contribution of walk, stop, and turn decisions in effective navigation, we systematically replaced each time-dependent rate with its average, so that overall biases were retained, but the dependency on encounters were not. Without encounter-modulated turning, adding stopping and walking decisions alone improved performance marginally (Figure 7D). With encounter-modulated turning present, however, the addition of either walk or stop decisions obeying our models both markedly increased the chance of finding the source and markedly reduced the search time. Together, this indicates a key benefit of encounter-driven stop-walk modulation when navigating spatiotemporally complex plumes.

**Figure 7:**
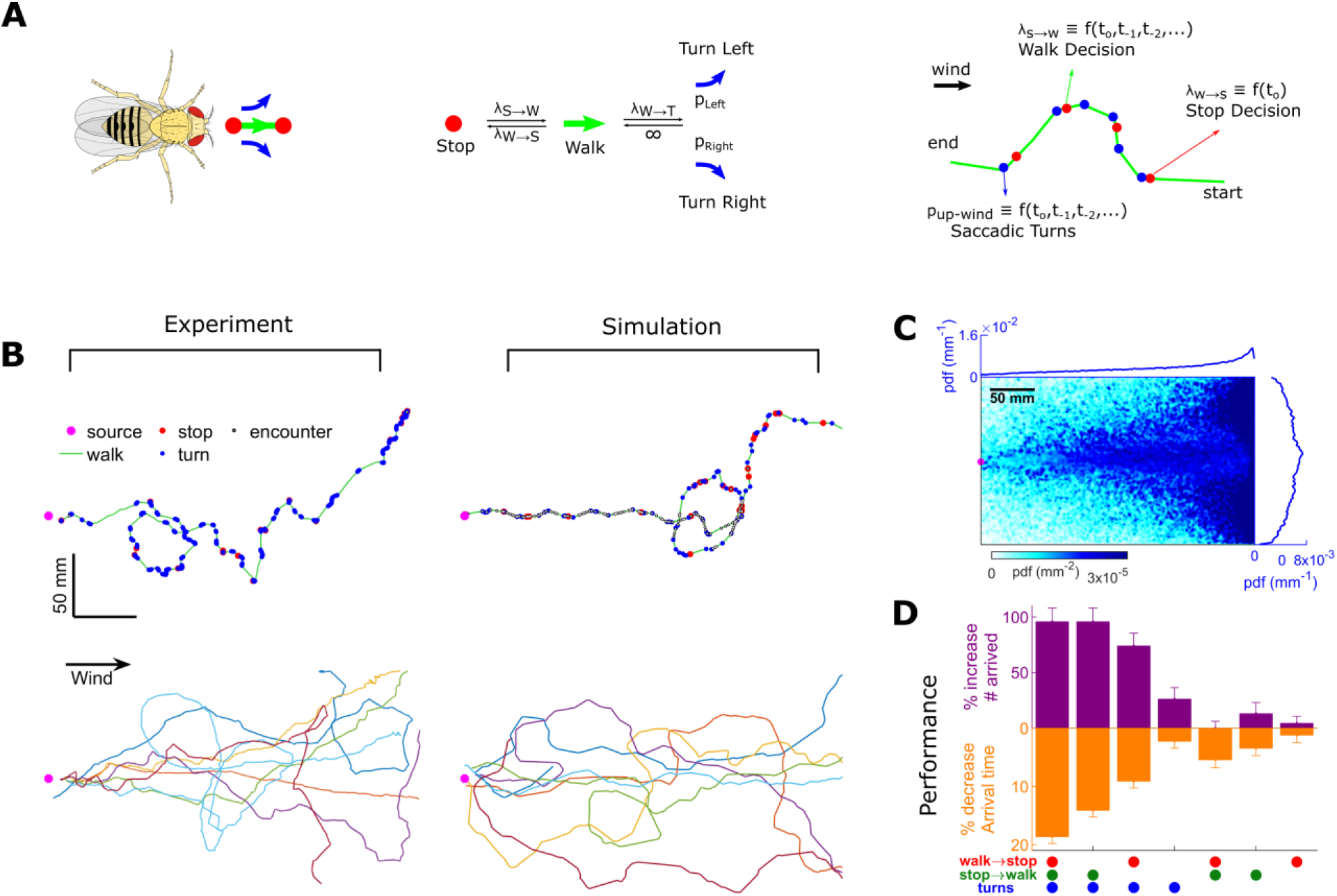
Agent-based simulation reveals navigational performance is significantly improved by encounter–modulated turn and walk decisions. **A.** Agent-based simulation. Left: An agent has four behavioral states: stops, walks, and saccadic turns either left or right. Middle: Diagram of behavioral transitions. Right: Hypothetical trajectory of a virtual fly. Stop-to-walk rates and upwind turn probability depend on encounter history, whereas walk-to-stop rates depends only on the time of the last encounter. **B.** Comparison of trajectories of real flies to those of virtual navigators in complex plume. Top row: Representative trajectories. Bottom row: 7 exemplary trajectories that reached within a 15 mm radius of the source. **C**. Pdf of virtual flies (*n*=10000 trajectories). Magenta: location of the source; blue curves: marginal pdfs over x- and y-direction. Virtual flies center toward the source (compare Figure 1C). **D**. Purple: fractional increase in the number of virtual flies that arrived within a 15 mm radius of the source, 100(*N_i_* — *N_c_*)/*N_c_*, where *N_C_* is the number of control flies that arrived to the source, and *N_i_* is the number of flies arrived to source for simulation condition *i*. Simulation conditions *i* (x-axis) are distinguished by which behavioral components (turn, walk, and stop decisions) were removed. Stop and walk decisions were removed by setting the corresponding transition rate to their averages over all navigators in the full model (left-most bar). Turn decisions were removed by setting the upwind turn probability to its average. Orange: fractional decrease in the arrival time, 100(*T_C_* – *T_i_*)/*T_c_*, where *T* is the time to arrive to source. *n*= 450, 450, 400, 290, 230, 260, 240 trajectories. Error bars represent the SEM calculated by bootstrapping the data 30 times with replacement.

## DISCUSSION

Odor plumes can vary widely in spatiotemporal structure depending on the geometry of the surroundings and the nature of the airflow. In turbulent flows, the duration of odor encounters and blanks are power-law distributed, spanning a wide range of values from milliseconds to a few seconds long [1]. While these flows become laminar near very smooth boundaries, the presence innatural terrains of obstacles, wind shifts, source motions, surface roughness, and boundary layer instabilities can cause smooth odor streams to break up into complex filaments [10, 23–25]. In our wind tunnel, we generate such perturbations by perturbing the laminar flow with stochastically alternating air jets near the upwind end. The key feature of this environment is that the statistics of the resulting odor patches are broadly distributed and approximate those in the atmospheric boundary layer (Figure 1G–1H), while allowing us to image behavior and signal simultaneously.

The intermittent nature of turbulent odor plumes has inspired a number of theoretical navigational algorithms that treat odor signals as a train of event times, (e.g. as in w(t)), ignoring encounter information about concentration and duration [31, 32]. Indeed, information-theoretic analysis has indicated that precise measurements of odor concentration may confer less benefit than coarse measurements across space or time [51]. In ‘infotactic’ searches [32], agents successfully navigate turbulent plumes by updating an internal spatial model of the plume structure, using only the arrival time of individual encounters. Analysis of insect flying trajectories [52] and *C. elegans* larvae crawling patterns [53] indicate that encounter-timing-driven infotaxis may form part of the navigation repertoire when concentration gradients are absent or difficult to measure.

Beyond theory, various experiments have shown that in intermittent plumes the frequency of encounters strongly shapes navigational behavior. The starkest indication of this in insect olfaction is the response of flying moths, *Cadra cautella* and *Heliothos Virescens* [9, 13, 17, 54], and walking moths, *Bombyx mori* [12], to pheromone plumes. In turbulent plumes and plumes pulsed at sufficient frequency, moths follow tight, narrow trajectories toward the source, whereas when the pulsing frequency is too low or the ribbon is static, they execute more zigzagging motion. To explain this, a model has been proposed in which an internal counterturning tendency is suppressed or reset by plume hits [14, 17]. A loose analogy could be made with our findings. Moths move crosswind and execute counterturns to find the plume, but once within the plume cone, high frequency odor encounters cause them to suppress counterturns and surge upwind. Analogously, walking *Drosophila* move crosswind and execute local search to get inside the plume, but once inside a complex plume cone they execute random left/right saccades, with frequent odor encounters biasing these saccades upwind. In both cases, the timing and frequency of odor encounters suppresses exploration and drives progress towards the source.

Like moths, flying flies navigating static odor ribbons counterturn back into them after passing through, effecting a similar upwind zigzag motion, though with smaller angles [18, 19]. An alternative explanation to the internal counterturning model in moth is that flies simply counterturn after losing the plume [19]. The duration of encounters as flying flies pass through a static ribbon are brief – 10-250 ms – not unlike the encounters we measure in the plume used here. Further, due to the erratic zigzags of flies as they cross the ribbon, encounters are perceived somewhat randomly in time. Thus, it was suggested that since the statistics of perceived odor signals end up resembling those in turbulence, this plume loss-initiated counterturning might be a generic navigational strategy, occurring in spatiotemporally complex plumes as well [19].

At least for walking flies, we find that this is not the case. Turns occur stochastically, with rates independent of how long flies spend in the odor and the frequency of encounters (Figure 4C). But there is an important distinction between intermittency in flies crossing standing ribbons and those navigating dynamic plumes. In ribbons, intermittency is generated by animals’ self-motion, creating a strong correlation between the likelihood of an odor encounter and spatial location. The location of expected plume encounters is in this sense highly predictable, which makes counterturning an effective strategy. Within the cone subtended by our dynamic plumes, the frequency and duration of encounters are less correlated with location and direction, and can occur even when the fly is stopped. This makes the location of future hits less predictable (Figure 1J), so within the plume cone, reactive strategies such as counterturning might be ineffective.

An important finding here is that the duration of odor encounters plays no role in navigation (Figure 3D-F). This was unexpected, since a recent systematic quantification of navigation algorithms in walking *Drosophila* found that flies bias their orientation upwind by integrating odor concentration [20]. In that model, the concentration is normalized, so reorientations are accounted for primarily by the duration of the odor. However, the odor signals were relatively slow – pulsed from 0.1 to 1 Hz – giving encounter durations an order of magnitude larger than in the plume used here. This suggests that for rapid, intermittent signals, encounter frequency drives navigation, while for slower signals such as those expected in the boundary layer of a smooth surface, the duration of odor exposure matters. Effective navigation may therefore combine two important features of the temporal odor signal: its rectified derivative (giving encounter onset times) and its integral (giving odor exposure time). Future studies interpolating between these extremes could elucidate if and how animals weight these two distinctly informative contributions.

A second important finding is that stopping forms a key component of the search strategy for walking flies (Figures 5–7). Stopping and waiting for encounters allows flies to receive odor encounters from dynamic plumes without wandering off-track or expending energy. We find that in deciding to walk, flies accumulate evidence from individual encounters, so walks are more likely following a clump of encounters than a single one. Theoretical work has shown that evidence accumulation from odor encounters can inform internal representations of plume structure to drive successful navigation in gradient-less plumes [32, 53], an interesting possibility still to be examined. Filtering and integrating odor concentration drives navigation in odor plumes with longer encounters and less regularity [20], suggesting that evidence accumulation – be it from odor duration or frequency – is a generic feature of olfactory navigation in a variety of environments. More work is required to understand the neural circuits and computations responsible for enacting stop and walk decisions. There is evidence that the transcription factor *FoxP* plays a role in value-based decision making, implicating these mutants as possible targets for future studies [55]. Finally, encounter-elicited stopping might be unique to walking *Drosophila* and larvae [56], since remaining stationary is more difficult in flight. Still, the reflexive counterturns that flying *Drosophila* execute after losing a plume [19] do bear loose resemblance to the increased stop rates following a drop in encounter frequency, so these decisions may have a common origin, but a different behavioral response.

It is surprising that despite the rich locomotive repertoire of walking *Drosophila*, a large part of their olfactory navigational strategy can be reduced to four actions – left turn, right turn, walk and stop. Still, we expect that differences in wind conditions and in odor identity and valence might modulate finer motor control in navigation [39]. A recent, systematic study of the locomotive structure of walking *Drosophila* in various odor environments without horizontal wind has found that behaviors fall into a limited number of states comprising a hierarchical hidden Markov model [57]. While the identity of the odor and fly individuality affect the transition rates between these states, new states do not emerge in different conditions. These findings are consistent with ours. A natural extension would be to study how fly individuality and odor identity affect transition rates in our model, and which conditions would indeed require an extended behavioral space.

Finally, an important aspect not explored in our work is learning. The navigational algorithms we have found in the plume used here are shaped by odor information from the recent past, over timescales no longer than a few seconds. Animals can learn odor landscapes over longer periods, by associating odor cues with location. Desert ants, *Cataglyphis fortis*, have been shown to use learned olfactory scenes for homeward navigation in the absence of other directional cues [58]. Similarly, in mice, efficient foraging strategies can overtake an otherwise local gradient ascent strategy, if prior information about the odor scene is available [59]. It is possible that the stochastic random walk strategies we observe here could be replaced with more stereotyped maneuvers if flies were sufficiently preconditioned to the environment. How the navigational strategies we have observed here are affected by conditioning, either with repeated trials or with reward feedback, provides a fruitful direction for future studies.

## ACKNOWLEDGEMENTS

We are thankful to John Carlson for providing several fly lines used in this study and for giving us access to instruments in his lab. We thank Richard Benton for providing anosmic flies, Francesco Carbone and Kevin Gleason for performing GCMS analysis on the smoke, Srinivas Gorur-Shandilya for help writing the fly tracking code, and Pedro Cisneros and Abhishek Sethi for their help writing scripts and annotating experimental videos. M.D. and T.E. were partially supported by the Allen Distinguished Investigator Program (grant 11562) through The Paul G. Allen Frontiers Group. T.E. was partially supported by NIH R01GM106189. N.K. was supported by a postdoctoral fellowship through the Swartz Foundation and by postdoctoral fellowship NIH F32MH118700. H.D.A. was supported by NSF DBI-1755494. D.A.C. was supported by NIH R01EY026555, a Searle Scholar Award, a Sloan Fellowship in Neuroscience, and the Smith Family Foundation.

## AUTHOR CONTRIBUTIONS

M.D. and N.K. contributed equally to this work. M.D. and T.E. conceived the project and designed the experiments with help from D.C. M.D. performed all experiments. N.K, M.D., D.C. and T.E. performed the data analysis. N.K. performed the mathematical modeling. H.A. performed the agent-based simulations. N.K., M.D. and T.E. wrote the manuscript. All authors edited the manuscript.

## MATERIALS AND METHODS

### Key resources

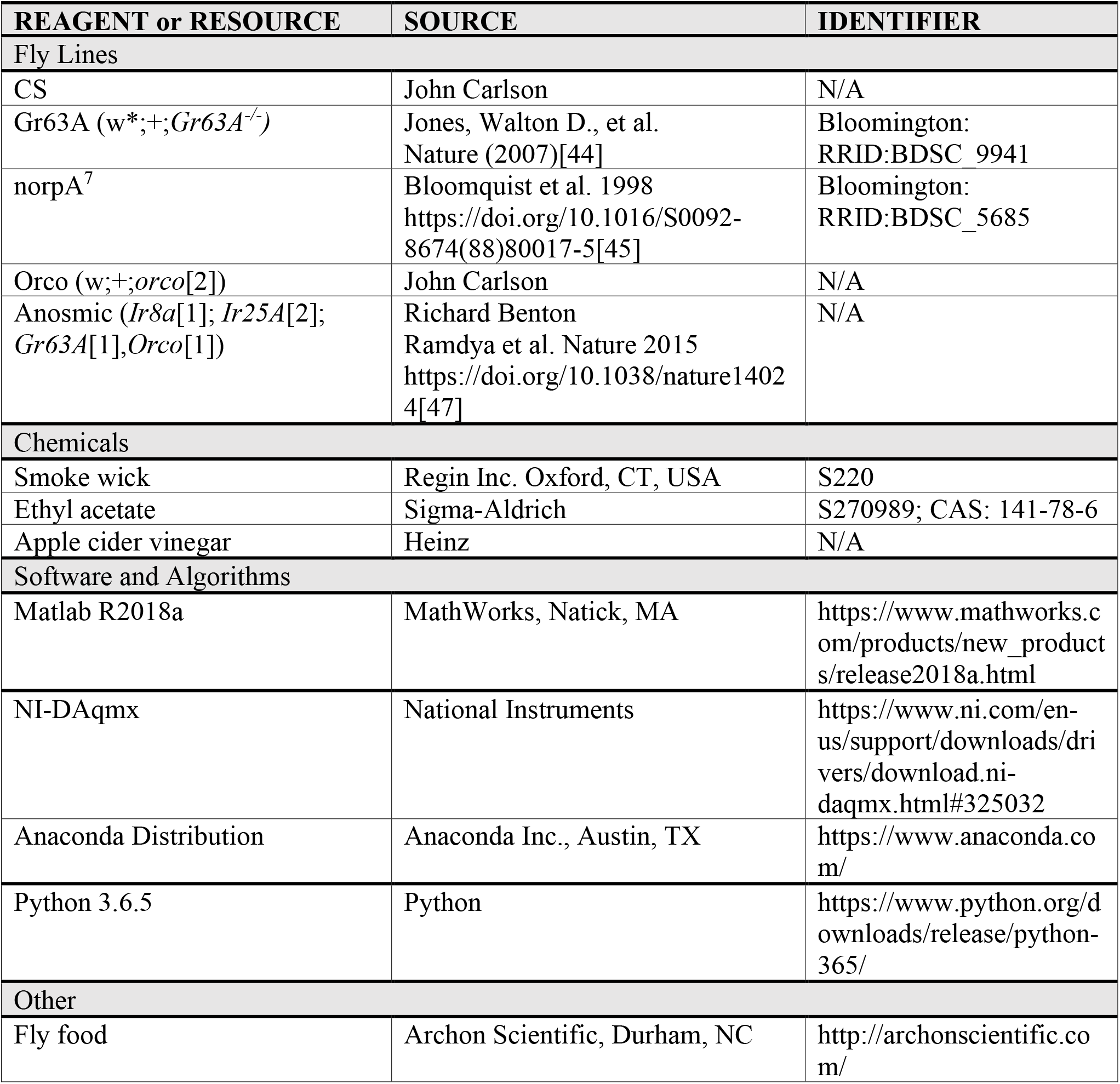

## LEAD CONTACT AND MATERIALS AVAILABILITY

Further information and request for reagents may be directed to, and will be fulfilled by the lead contact

## EXPERIMENTAL DETAILS

### Fly strains and handling

Flies were reared at 25°C and 60% humidity on a 12h/12h light-dark cycle in plastic vials containing 10 ml standard glucose-cornmeal medium (i.e. 81.8% water, 00.6% agar, 05.3% cornmeal, 03.8% yeast, 07.6% glucose, 00.5% propionic acid, 00.1% methylparaben, and 00.3% ethanol. Media was supplied by Archon Scientific, NC). All flies used in behavioral experiments were females. Newly eclosed flies were collected each day and placed in fresh vials. Females were then collected for starvation and placed in empty vials, 30-40 females in each vial, containing soaked cotton plugs at the bottom and top. All flies were 5-9 days old and 3-4 days starved when experiments were performed. Experiments were carried out within 5 hours prior to the subjective sunset (i.e. 12hr light turn off). All fly strains used in this paper are listed in Key Resources Table.

### Behavioral apparatus and stimulus delivery

Flies are introduced into an arena of size 300 mm (along wind) and 180 mm (across wind) with a depth of 10 mm (see schematic, Figure 1A). Flies walk unrestrained in this arena on glass surfaces, top and bottom, which were separated with acrylic walls and also bounded at the upstream and downstream end of the arena by straws and a plastic mesh, respectively. Experiments were recorded with an infrared (IR) sensitive camera (FLIR Grasshopper USB 3.0 NIR) in a dark room under IR illumination (850 nm). The recording rate was 30 Hz and 90 Hz for straight and intermittent plume experiments respectively. The intermittent plume required a higher frame rate to track the dynamic smoke stimulus with sufficient resolution.

The behavioral apparatus operates as a wind tunnel. Active-charcoal filtered dry-air passes through the straws stacked at the upstream end of the arena and creates a laminar flow with a flow speed around 150 mm/sec. Flow speed is measured by two methods: with an anemometer and by imaging and calculating the speed of a smoke plume in the laminar flow. The flow speed calculated with these two methods were similar (data not shown). The air and any odor carried with it were collected at the downstream end of the device with a vacuum hose loosely coupled to the device.

In order to deliver odorants in to the behavioral chamber, clean air was passed over the headspace of pure odors placed in glass vials, and obtained odorized air was passed through a straw fixed at the center of the stack that creates the laminar flow. Variations in the odor dose was obtained by varying the ratio of odorized and clean air in the final flow delivered in to the chamber. In the case of smoke, smoke generated by a burned wick (S220, Regin Inc.) is accumulated in a 250 ml bottle for 20 seconds, and that bottle was used as the smoke-odor source. Pure ethyl acetate was purchased from Sigma-Aldrich and apple cider vinegar was made by Heinz.

Straight plumes were obtained by simply matching the odorized air flow speed to laminar flow speed. In order to generate intermittent plumes, air flows (flow speed: ~1500 mm/sec) perpendicular to the laminar flow were injected in to the arena near the upstream end of the device. The injected air flows were randomly alternated between left and right side of the arena with 100 ms correlation time.

### Experimental protocol

Starved flies, between 30 and 40 in number, were aspirated in to the arena all in once while wind was on, and allowed 1 min to acclimate to their environment prior to the experiment. Odor was on during the whole experiment, which lasted 90 seconds (unless otherwise noted), however it took several seconds, ~5s, to pass the tubing and enter to the arena. In pulsed intermittent plume experiments, while the random lateral air injections persisted, odor was alternatively turned on and off in 15-second blocks. Experiments were repeated 3 times on the same flies, leaving 5 min intervals in between. Glass surfaces are wipe cleaned with Windex before each experiment, and the whole stack of straws and tubing are replaced with clean ones before a different odor is tested. The humidity and temperature of the room is logged for each experiment. Room temperature was stable around 24.2 ± 0.3 °C. Although the room humidity varied between 25% and 43% (average: 32.5 ± 2.5%) depending on the season of the year, flies were tested in dry air flow which had humidity close to zero.

## QUANTIFICATION AND STATISTICAL ANALYSIS

All analysis was performed using custom written Matlab and Python scripts.

### Fly tracking and signal estimation

#### Fly tracking

Fly tracks were extracted by analyzing recorded videos. Non-uniform illumination in the arena was corrected by diving each frame pixel-by-pixel with a flat field image. The flat field image was obtained as follows: a) median smoothing (filter size ~ 8mm) the image of the arena free of flies, b) fitting a 5^th^ order polynomial to the smoothed surface, c) normalizing the fitted surface with the mean value. Following the flat-field correction, each frame was thresholded and binarized. In the binarized images, objects with an area larger than one square millimeter were registered as flies. The positions (*x* and *y* coordinates) and orientations (*θ*) of the flies were obtained by finding the centroids and major axis of these regions (Matlab regionprops). Tracks were assembled by linking each centroid in a frame to the closest one in the consecutive frame. Whenever two or more flies interacted (i.e. passing over each other on opposite surfaces, or collision), water-shedding was used to resolve the identity of the flies. If water-shedding failed, fly positions were predicted based on their recent average velocities until they separate, and identities were then assigned by comparing the predicted positions with the positions right after separation. If this failed the track was flagged and eliminated from subsequent analysis.

#### Signal estimation

The signal *s*(*t*) that flies experience along their path was estimated by calculating the mean smoke intensity in a virtual antenna fixed in front of the flies’ head (Figure 1B inset). The virtual antenna was as wide as the fly (i.e. 1.72±.24 mm), and its length was set to 1/5^th^ of the fly minor axis (average: 0.46±.08 mm). The distance between the virtual antenna and fly head (i.e. 1.24±.22 mm) was optimized by minimizing the overlap between the virtual antenna and dilated fly body (dilation number: 7). Instances in which the virtual antenna overlapped with another fly or its reflection were flagged as unreliable and eliminated from all analyses that required odor information. The accuracy of our automated detection system was validated by comparing its output to the output of manual annotation. 4 videos were manually annotated, frame by frame, by two researchers for validity of the signal values in the virtual antenna of all flies. The total number of annotated time points was 166411. This comparison revealed that, only 0.97% of all data points were assigned as false positives by our automated software, whereas the fraction of trajectories that were assigned as false negatives was 6.05%.

#### Encounter detection

Due to shot noise, signal values were above zero. In order to calculate the mean background signal value, for each recorded video, we fitted a Gaussian to the distribution of the signal values of all flies, and set the mean signal background to the mean of the Gaussian. The signal was then thresholded with a threshold value, equal to the sum of the mean and 2.5 SD of the background signal, and binarized for encounter detections. Instances with signal values higher than the threshold are identified as encounters, and blanks are defined as the intervals between encounters.

#### Error estimation in the timing of encounter detection

The difference in the location of the virtual antenna from that of the actual fly antenna introduces some uncertainty in signal timing. This uncertainty depends on wind speed and direction, and we estimated it as 8.3±1.5 ms and −8.3±1.5 ms for downwind and upwind oriented flies, respectively. Further uncertainty in timing is introduced by our encounter threshold, which we chose as 2.5 standard deviations above the mean camera shot noise. To estimate this uncertainty, we considered flies navigating a static smoke ribbon. We calculated the lateral distance from the plume at which flies counterturn back into the ribbon after passing through it (counterturn positions were found using Matlab’s “findpeaks” function), and compared it to the iso-line of the smoke intensity used for the encounter threshold (2.5σ) (Figure S2A). The average counterturn positions along the ribbon (calculated using Matlab’s LOWESS smoothing function) aligned with the iso-line of the minimum smoke intensity captured by the camera (Figure S2B). However, they differed from the iso-line of our 2.5σ encounter threshold by 1.94 mm at the downstream end of the device, giving an upper bound of 13 ms uncertainty in encounter timing. Combined with uncertainty from the virtual antenna locations, we estimate the overall error in encounter timing to be less than 25 ms.

### Analysis of behavioral data

#### Smoothing of measured behavioral time traces

From the procedures described above, we obtained for each fly trajectory the position (*x* and *y* coordinates) and the orientation (*θ*) of the fly, together with the signal *s* in the virtual antenna. To remove measurement shot noise from the fly tracking and signal estimation, we filtered each of these quantities with a Savitsky-Golay filter with *k*-order polynomial and *m*-length windows, where *k* = 4 and *m* = 9. Taking the derivative of the fitted piecewise Savitsky-Golay polynomials for *x* and *y* gives us smoothed velocity components and *v_y_*, respectively, from which we obtain the speed 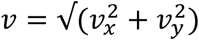. Similarly, the time rate of change of orientation, *ω* = *dθ*/*dt*, was found by converting *θ* to *x-y* components on the unit circle, smoothing each of these with the same Savitsky-Golay filter, taking their first derivative (which is the analytical derivative of the smoothed polynomials), and then converting back to polar coordinates. This last conversion was done using 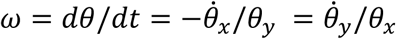, the appropriate equation being chosen if either *θ_x_* or *θ_y_* were too small. This procedure removes issues with the branch cut at *θ* = 0, which would arise if *θ* were differentiated directly. Finally, upwind orientation *θ*_+_ was determined by reflecting all angles over the polar x-axis to the range 0-180 degrees. All subsequent analyses used these smoothed time traces.

#### Determination of stop, turn, and encounter events

From these smoothed quantities, all binary events – stops, turns, and encounters – were determined in the same way, using a thresholding technique that minimizes false detections. Specifically, the onset of an event is said to occur when the quantity is above threshold, but only if the time above the threshold is longer than some set duration. This prevents false detections of artificially short events that may arise from measurement fluctuations. The same requirement is enforced for the event offset: the quantity must drop below threshold for a sufficient time. For encounter instances, the threshold was set to 2.5X the standard deviation of the background signal, when fit to a Gaussian. For stops, the threshold was set at *ν* = 2 mm/s, and for turns the threshold was set to |*ω*| = 200 deg/s. The minimum duration for stops was set to 300 ms, for turns was set to 20 ms, and for encounters was set to 50 ms.

To verify that the discrete nature of turns was not an artifact of turn detection thresholds, we used the following analysis, whose results are presented in Figure S5. Turn events were first detected for a given threshold on absolute angular velocity, |*ω*|. A window of length Δ*T* is defined around the midpoint of each turn, over which the total orientation change Δ*θ* of the fly can be defined (Figure S5A, top plot). If angle changes are indeed discrete, then the distribution of Δ*θ* would increase with Δ*T* for small Δ*T*, but would level out asymptotically to a bimodal distribution for sufficient Δ*T*, corresponding to the maximum length of a stereotypical turn. For different choices of |*ω*| (different colored plots in Figure S5B), darker shades correspond to Δ*θ* distributions for increasing window lengths Δ*T*. The distributions are invariant once the window is long enough, showing that for sufficiently long time window, the turns are discrete. The positive peak of the distributions is plotted as a function of Δ*T* (Figure S5C). This peak levels off above Δ*T* = 150 ms, for all turn thresholds larger than 150 deg/s. As such, we chose a threshold of 200 deg/s for turns, verifying visually that false positives and false negatives were minimized.

#### Fly distributions in arena

The probability density function (pdf) of fly distribution in the arena (Figures 1K–1L, 7C, S1C, S1E, S1G, and S1I) is estimated by calculating the histogram (count) of fly positions for each unit area (ΔX= ΔY=1 mm), and normalizing that histogram with the total count of fly positions. Areas with zero fly visits are indicated with white color in the color scale. All positions for all time points throughout the experiment is used to calculate the histogram, and therefore estimated pdf represents the cumulative fly walking behavior.

#### Calculation of stop-to-walk and walk-to-stop transition rates

To calculate the transition rate from a stop to a walk *λ_s→w_* (Figure 2F), we considered each part of the trajectory containing one stop followed by one walk. In this range, we set 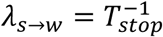, where *T_stop_* is the duration of the stop. Likewise, we get the transition rate from walk to stop, *λ_w→s_*, by considering every trajectory snippet containing one walk followed by one stop. In this range, 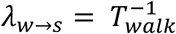. Thus, the walk-to-stop rates update their values at the onset of every stop, and vice versa for walks.

### Numerical methods

#### Statistical tests

Unless noted otherwise, all error bars represent standard error of the mean. Stars in the manuscript indicating significance are * (*p*< 0.05), ** (*p*< 0.01), *** (*p* < 0.001). Unless otherwise noted, for estimating SEM through bootstrapping, 5000 resamples were chosen, with resample size half of the original data size. The trivariate linear regression we use to determine the dependence of upwind orientation *θ*_+_ on *W_dur_*, *W_freq_*, and *W_conc_* (Figure 3F) can exhibit issues of multicollinearity when the two regressors are correlated. This can produce erroneous and non-robust parameter estimates. To check for this, we ensured that the moment matrix *X^T^X*, where *X* is the matrix of observations, was not ill-conditioned. We found conditions numbers less than 2, indicating no ill-conditioning. We also ensured that the estimated parameters were robust to different subsets of the data. We found that the parameter ranges for these subsets are statistically indistinct from those of the full dataset.

#### Calculation of encounter frequency and encounter duration

Calculation of the odor encounter frequency *W_freq_* was done by convolving the binary vector of encounter times *w*(*t*), which contains a ‘1’ at the onset of every encounter and 0s elsewhere, with an exponential filter:

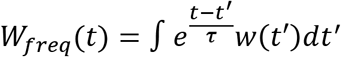

where *τ* is set to 2 seconds. This timescale was chosen to optimize the trends, but our findings were robust to this value. Similarly, the encounter duration *W_dur_* was calculated by convolving the binary vector of encounter exposure, *d*(*t*) which contains a ‘1’ whenever the fly is within a encounter and 0s elsewhere, with the same exponential filter:

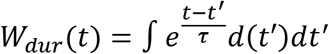

Finally, odor concentration was calculated by convolving the raw signal *s*(*t*) (an 8-bit integer representing the intensity of the imaged smoke signal) with the same exponential filter:

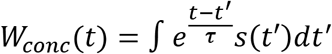

### Modeling

#### Stochastic models of stereotyped turning

Turns are modeled as a homogeneous Poisson processes with rate 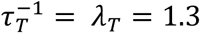 turns/s, determined by fitting the data to an exponential distribution (Figure 4C). While we did not find a statistically significant dependence of the turning rate on encounter frequency or duration, we allowed for this dependence in fitting the model to the data (below). When a turn occurs, its magnitude and direction are also chosen from random distributions. The absolute value of the turn angle is chosen from a normal distribution with mean 30° and standard deviation 10°, representing the two peaks in the bivariate distribution in Figure 4A. The turn direction is a binomial variable with probability *p_T_*(*w*(*t*)) that the direction is upwind, where the encounter onset times *w*(*t*) is defined above. Guided by the observation that the upwind turning probability increases with encounter frequency, we set *p_T_*(*w*(*t*)) as:

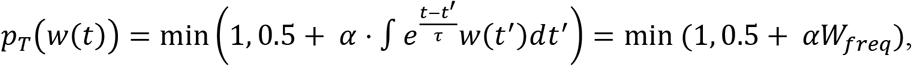

where the filter timescale *τ* is set to 2 seconds as above. In this model, the likelihood that turns are upwind increases by *α* with each encounter, and decays to 0.5 otherwise. The “min” is taken to keep the probabilities bounded between 0 and 1.

#### Stochastic accumulated evidence,” “last encounter, “ and “odor duration” models of walk-stop transitions

Walk-to-stop transitions are modeled as an inhomogeneous Poisson process with rate *λ_w→s_*(*t*) = *λ_w→s_*(*w*(*t*), *d*(*t*)), where *w*(*t*) and *d*(*t*) are defined above. The distinguishing feature between the three models we test, the last encounter model, the accumulated evidence model, and the encounter duration model, is their functional dependence on *w*(*t*) and *d*(*t*).

In the last encounter stopping model, we have:

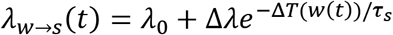

where Δ*T*(*w*(*t*)) indicates the time since the most recent encounter. In the accumulated evidence stopping model we have:

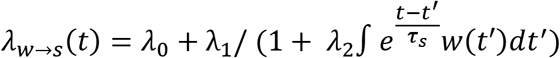

Finally, in the encounter duration model, we have:

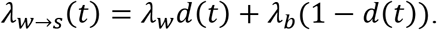

In the encounter duration model, therefore, the rate is *λ_w_* during encounters and *λ_b_* during blanks. The rates for the stop-to-walk transitions are analogous. For the last encounter model we have:

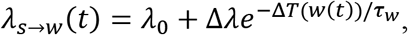

for the accumulated evidence walking model we have:

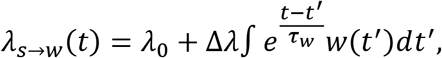

and for the encounter duration model we have:

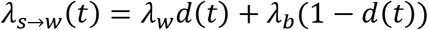

#### Turn model parameter estimation

In our turning model, the gain parameter *α* must be estimated from data. While we measured the turn rate *τ_T_* from experiment and found no significant dependence on encounter duration or frequency (Figure 4C), we nevertheless allow it to be a free parameter with linear dependence on both. The full turn rate 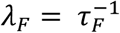 is therefore:

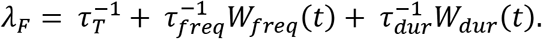

The turning model thus contains 4 unknown parameters: Θ = *α, τ_T_, τ_freq_, τ_dur_*, which we obtain using maximum likelihood estimation. The expression for the maximum likelihood depends on various conditional probability distributions, which we now discuss. The measured data are orientation changes *dθ_i_* at each time step during which the fly is walking. These angle changes arise either during straight bouts, where the angle changes consist of small zero-mean jitter, or during turns. At any given time *i*, the probability of angle change *dθ_i_* is:

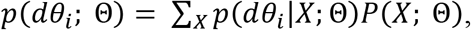

where *X* denotes the 3 possible behaviors: upwind turns, downwind turns, and straight bouts. The probability that a turn is upwind, *p*(*up*; Θ), is the probability of a turn times the probability that the turn is upwind, given by our turn model above. This gives

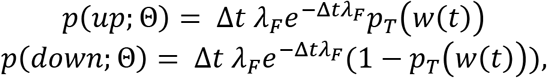

where Δ*t* is time step of the measured data. The distribution of upwind angle changes, *p*(*dθ_i_*|*up*; Θ), is assumed Gaussian with *σ* = 10° and mean +30° (for *θ_i_* between 0 and 180°) and —30° (for *θ_i_* between 180° and 360°). These values represent the mean and spread of the two peaks in the measured *dθ_i_* during turns (Figure 4A). The distribution of downwind angle changes *p*(*dθ_i_*|*down*; Θ) is the same, but with opposite means. The likelihood of a straight walk is:

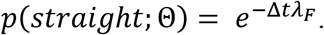

The jitter for straight bouts was measured to be 0.22° per time step, whereby *p*(*dθ_i_* |*straight*; Θ) is assumed normal with standard deviation 0.22° and mean θ. With these distributions, the likelihood function of parameters Θ given the measured data *dθ_i_* reads:

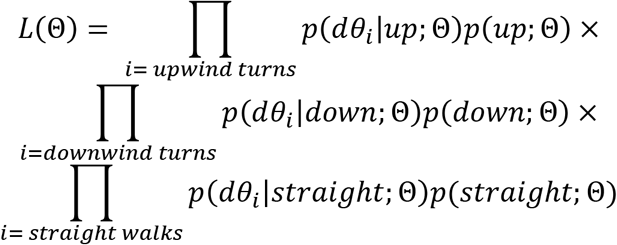

The first product contains only times at which the flies turn upwind, etc.. The parameters were estimated by maximizing the logarithm of *L*(Θ). The optimization was performed using the limited-memory Broyden–Fletcher–Goldfarb–Shanno optimization algorithm (L-BFGS). In our implementation of this method, we only provide the cost function (the log-likelihood), and the gradient is computed numerically with finite differences. L-BGFS approximates Hessians from function and gradient evaluations in previous iterations, using this to steer the estimate toward the function minimum. To get an idea of the spread and robustness of parameter values, we perform this optimization for 500 distinct subsets of the data, where each subset contains 20% of all measured trajectories, randomly chosen. We find that the distribution of the 500 estimated *α* and *τ_T_* are highly peaked around a mean of 0.046Hz^-1^ and 0.75s, respectively, suggesting that these values are robust to various subsets of the experimental data. The estimated coefficients τ_freq_ and τ_dur_ are small (medians 0.008 and 0.07, respectively, which would each contribute <3% to the base timescale τ_T_, given the encounter statistics in our intermittent plume), and span both negative and positive values, suggesting that they are not robust. These are in accordance with the lack of dependence in turn frequency on encounter duration or frequency (Figure 4C).

#### Stop and walk models parameter estimation

The likelihood function for stop and walk models is similar to the turn model. Since the walk-to-stop and stop-to-walk rates are distinct inhomogeneous Poisson processes, with distinct functional forms *λ_s→w_*(*t*) and *λ_w→s_*(*t*) for the rates, respectively, we write the likelihood function for walk-to-stop transitions *L_W→S_*(Θ) only here. The other, *L_S→W_*(Θ), is analogous.

The likelihood function for walk-to-stop transitions is:

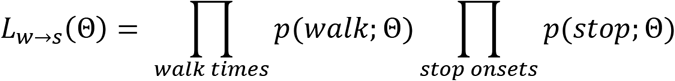

The first product runs over all times that the fly is walking, while the second product contains only the initial point of each stop bout, which represents the decision to stop, given that the fly is walking. Like the turning model, the rates follow a Poisson process, giving:

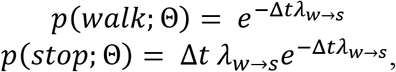

where the dependence on the parameters Θ enters through *λ_w→s_*. Parameters are estimated as described above, by minimizing the logarithm of the likelihood. Rather than optimizing for *τ_w_* or *τ_s_*, both of which enter the cost function in the denominators (introducing artificial singularities in the derivatives), we optimize for their inverses, which enter linearly in the exponents.

#### Generation of turn model predictions

To generate the upwind orientation versus encounter frequency plot predicted by the model estimates, (red line in second plot of Figure 4G), we chose from the 500 sets of estimated parameters Θ the median of each parameter. Synthetic *θ*(*t*) traces were generated by applying the model to a synthetic random encounter trace which contained encounter frequencies spanning a range from very low (< 0.1 Hz) to high (10 Hz), mimicking the range of frequencies encountered in the data.

#### Generation of stop and walk model statistics

To generate the statistics in Figures 5–6, we chose from the 500 sets of Θ the median of each parameter. Synthetic stops and walks were then generated from the aggregated measured *w*(*t*) of all trajectories by simulating a Poisson process with the corresponding rate functions. This procedure generated a synthetic time series of stop events, given that the fly is walking. From this vector, we calculated time-to-stop for all cases as in the measured data: i) all encounters, ii) all isolated encounters, and iii) all isolated clumps. A similar procedure was used to generate a vector of walk events, given that the fly is stopped. From this vector, we calculated time-to-walk for analogous cases: i) all encounters, ii) all isolated encounters, and iii) all isolated clumps.

### Agent-based simulation

Virtual agents navigated the same arena and the same intermittent smoke environment as experimental flies. Each simulation batch was run using 10,000 agents, initialized randomly along the back, downwind wall of the arena. The simulation ran for 8,000 time steps at a timestep corresponding to the same sampling rate as the recorded videos, 11.1 ms. The agents were assigned a walking speed equal to the mean walking speed of experimental flies (11.5 mm/s), and their turns were assumed to be instantaneous. Turn angles were drawn from a normal distribution with mean 30° and standard deviation 10°. The navigators were given elliptical virtual antennas, the centers of which were located approximately 14 pixels (2.16 mm) in front of the fly centers. The elliptical antennas had a semi-major axis of 5 pixels (0.77 mm) and semi-minor axis of 1.5 pixels (0.23 mm); the semi-major axis was oriented perpendicular to the agent’s orientation vector. To avoid spurious detections arising from jitter in the signal, the virtual navigators registered a encounter only if they had not encountered one in the past 100 ms.

To remove individual navigational components, the corresponding rate for that component was set to an average rate from the full model. For example, to remove stop decisions, the walk-to-stop rate was set to the average. The average stop-to-walk rate was determined by fitting the distribution of stop durations from all 10,000 trajectories in the full model to an exponential distribution; the fitted timescale was used as the average rate. The average walk-to-stop rate were determined similarly from the distribution of walk durations. The average probability of turning upwind was defined as the fraction of turns directed upwind by all 10,000 trajectories in the full model.

## DATA AND CODE AVAILABILITY

The data, fly lines used in this study and the scripts used to perform experiments, track flies and extract relevant behavioral data are available upon request. We thank the following people for making their Matlab scripts, utilized for generating plots in this work, freely available: Ben Mitch, Panel; Kelly Kearney, legendflex; Yair Altman, export_fig; David Legland, geom2D; Rob Campbell, shadedErrorBar.

## SUPPLEMENTAL INFORMATION

**Figure S1:**
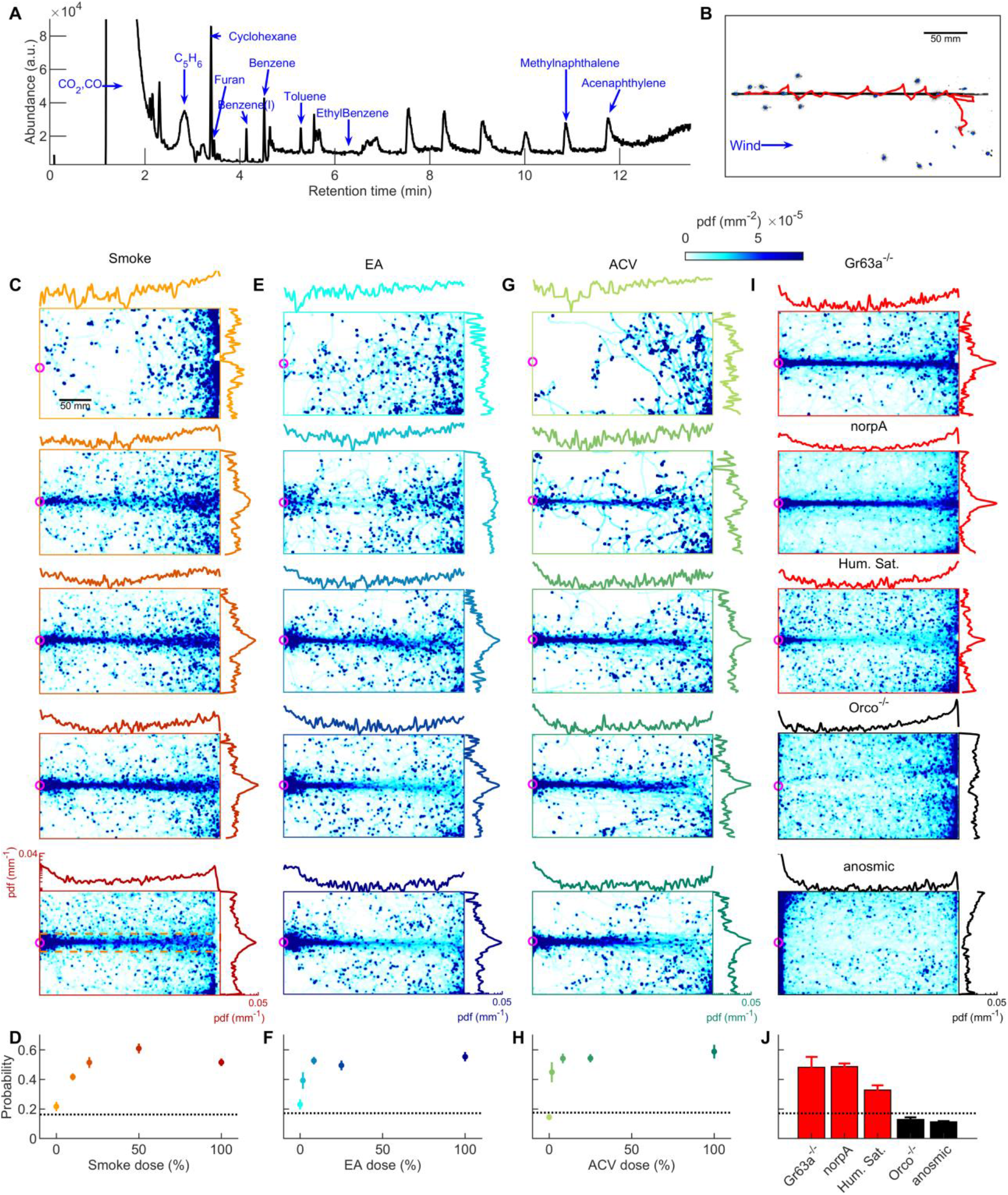
Attraction to smoke is olfactory and dose-dependent, closely mimicking that to ethyl acetate and apple cider vinegar. **A.** GC-MS measurement of smoke. Many volatiles belonging to various chemical groups such as, alcohols, ketones, and aldehydes are present in the smoke. **B**. A snapshot of flies (blue), navigating straight smoke plume (grey) in laminar flow (blue arrow, lateral jets were turned off) with a representative trajectory of a navigating fly (red). **C,** Probability distribution functions (pdf) of fly positions in the arena for wild-type CS flies navigating straight plumes (as in B) of smoke with increasing doses (top to bottom, *n*=871,1714,1600,1996,4421 trajectories). Marginal pdfs in the x- or y-direction are plotted on a log scale on the exterior. **D.** Integral of pdfs over the ribbon region (orange box, illustrated in the bottom plot of C) as a function of smoke dose. Dashed line represents chance probability. **E-F**. same as C-D but for ethyl acetate (EA, *n*=369,363,907,1351,1604), and **G-H.** same as C-D but for apple cider vinegar (ACV, *n*=176,212,829,1067,1332). **I.** Pdfs in straight smoke plumes for mutant flies, and at humidity saturation for wild-type CS flies. Mutant flies: Gr63a^-/-^ (*n*=1581); norpA (*n*=4480); Orco^-/-^ (*n*=2420); anosmic (Gr63a^-/-^, Orco^-/-^, Ir8a^-/-^, Ir25a^-/-^) (*n*=3992). High humidity: (86.0±3.9%) (*n*=2772). **J.** Pdf integrated over the ribbon region (same region used in F-H) for control experiments.

**Figure S2:**
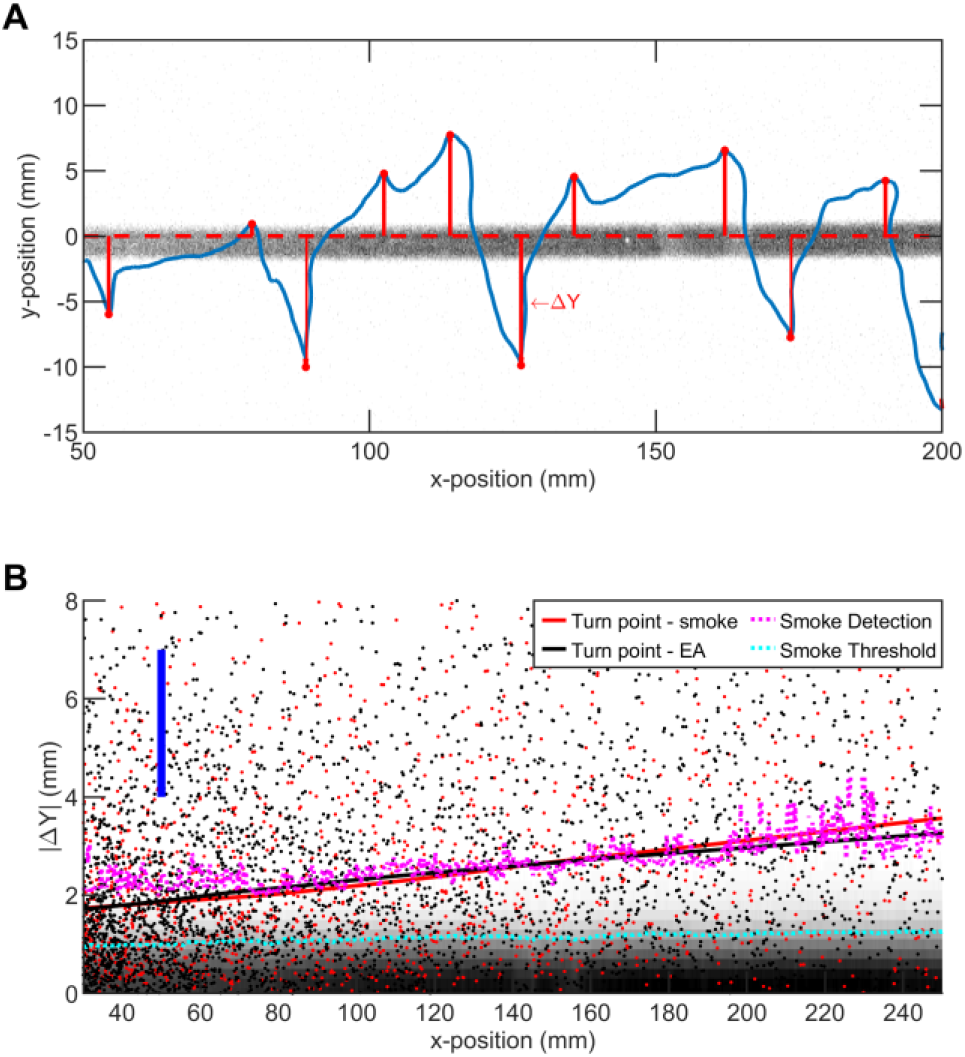
Average turning position of flies in straight plume agrees with smoke intensity. **A.** A representative trajectory of a fly walking around straight smoke plume or ribbon (lateral jets were turned off). Gray: smoke intensity. Blue: trajectory of the fly. Red Dots: points where fly makes a turn towards ribbon. Red Line: distance between ribbon center and turning point (i.e. ΔY) **B.** Comparison of smoke diffusivity to turning points of flies in straight plumes of smoke and EA. Red (Black) Dots: Turning points in straight smoke (EA) plume. Red (Black) curves: data smoothed with lowess method, with the option of resistance to the outliers. Blue bar: Length representing size of the fly (3mm) as a guide for the reader. Pink curve: iso-line of smoke intensity at 0.1 (a.u.). Cyan curve: iso-line of smoke intensity at 5 (a.u.) which is the value used for encounter detection.

**Figure S3:**
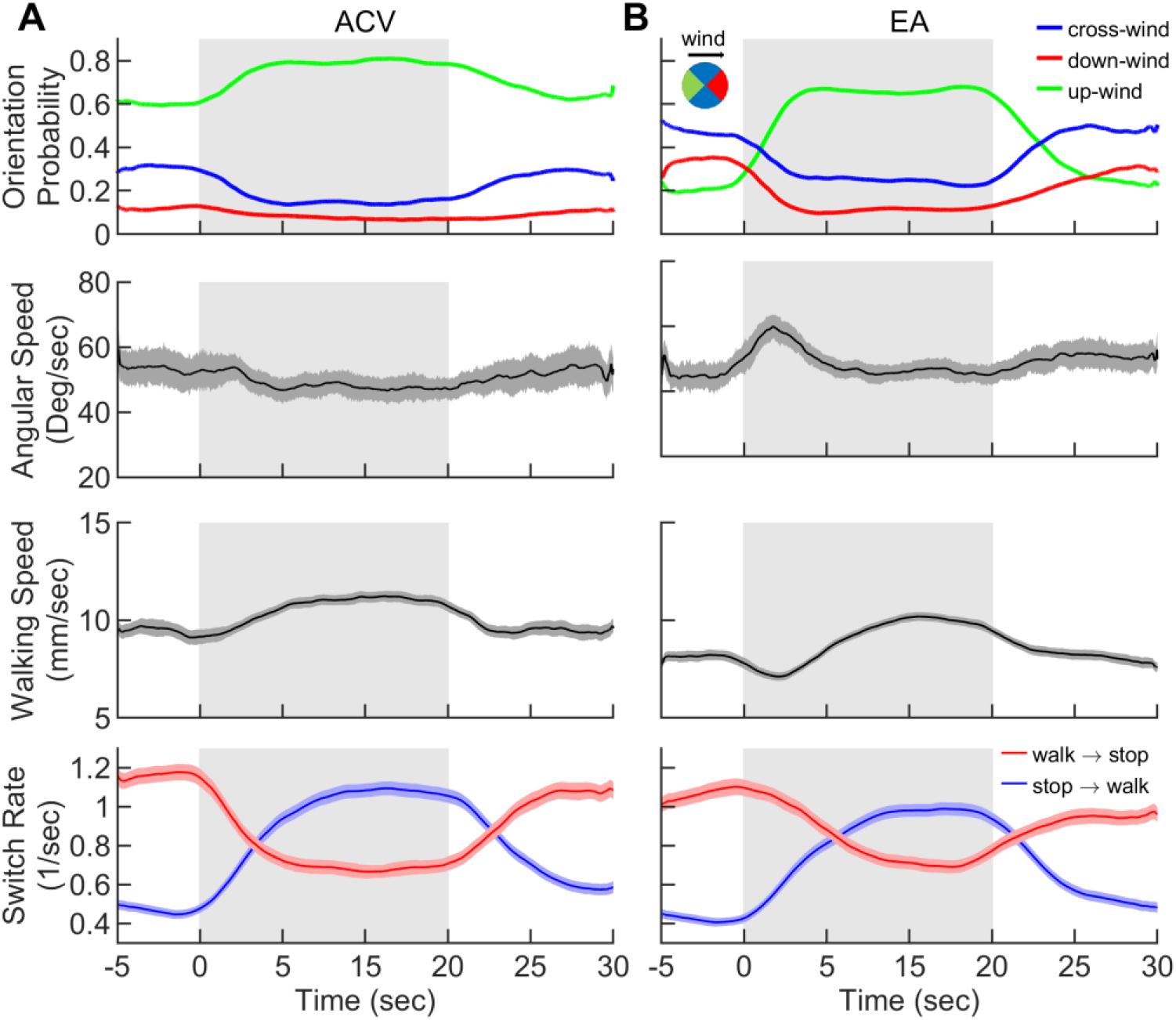
Smoke elicits behavior similar to natural odorants. Behavior of flies navigating in the intermittent plumes of **A.** apple cider vinegar (ACV) and **B.** ethyl acetate (EA) as in Figure 2 where wind is perturbed continuously with random lateral jets whereas the odor is turned on/off over 15 sec periods. Quantities averaged over all trajectories in A-B (n=1830 trajectories for ACV, and n=1809 for EA) as a function of time. **1^st^ row:** Probability of being in up-wind (green), cross-wind (blue) and down-wind (red) orientations estimated in 90 degree quadrants as shown in the circle with the same color codes. **2^nd^ row:** Angular speed **3^rd^ row:** Walking speed **4^th^ row:** Walk-to-stop (red) and stop-to-walk (blue) switching rates. Quantities in all four rows are smoothed with a 5-second sliding box filter. Error bars indicate SEM. Gray area indicates the period that odor injection, in to the arena, is turned while wind is constantly perturbed with lateral jets as in Figure 2.

**Figure S4:**
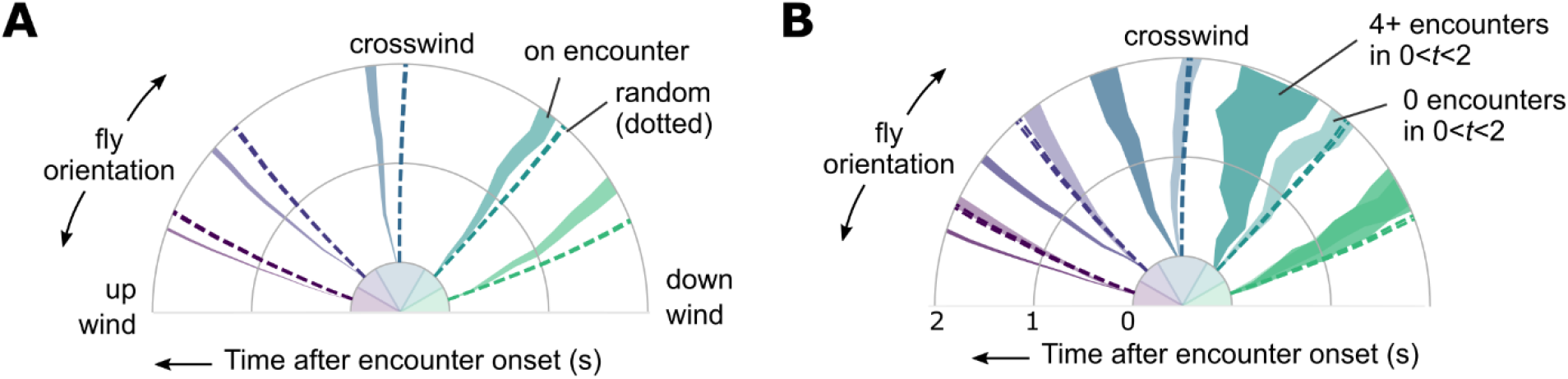
Encounter-elicited orientation change in time. **A.** Orientation 2 second window following an encounter (*n*=5040), for fly orientations binned in 0-30, 30-60, 60-120, 120-150, and 150-180 degrees, where 0 is upwind, and 180 is downwind. Time progresses radially outward. Orientations for random (non-encounter elicited) times are indicated by the dashed lines. **B**. Same data, now binned by number of subsequent encounters in the 2s window.

**Figure S5:**
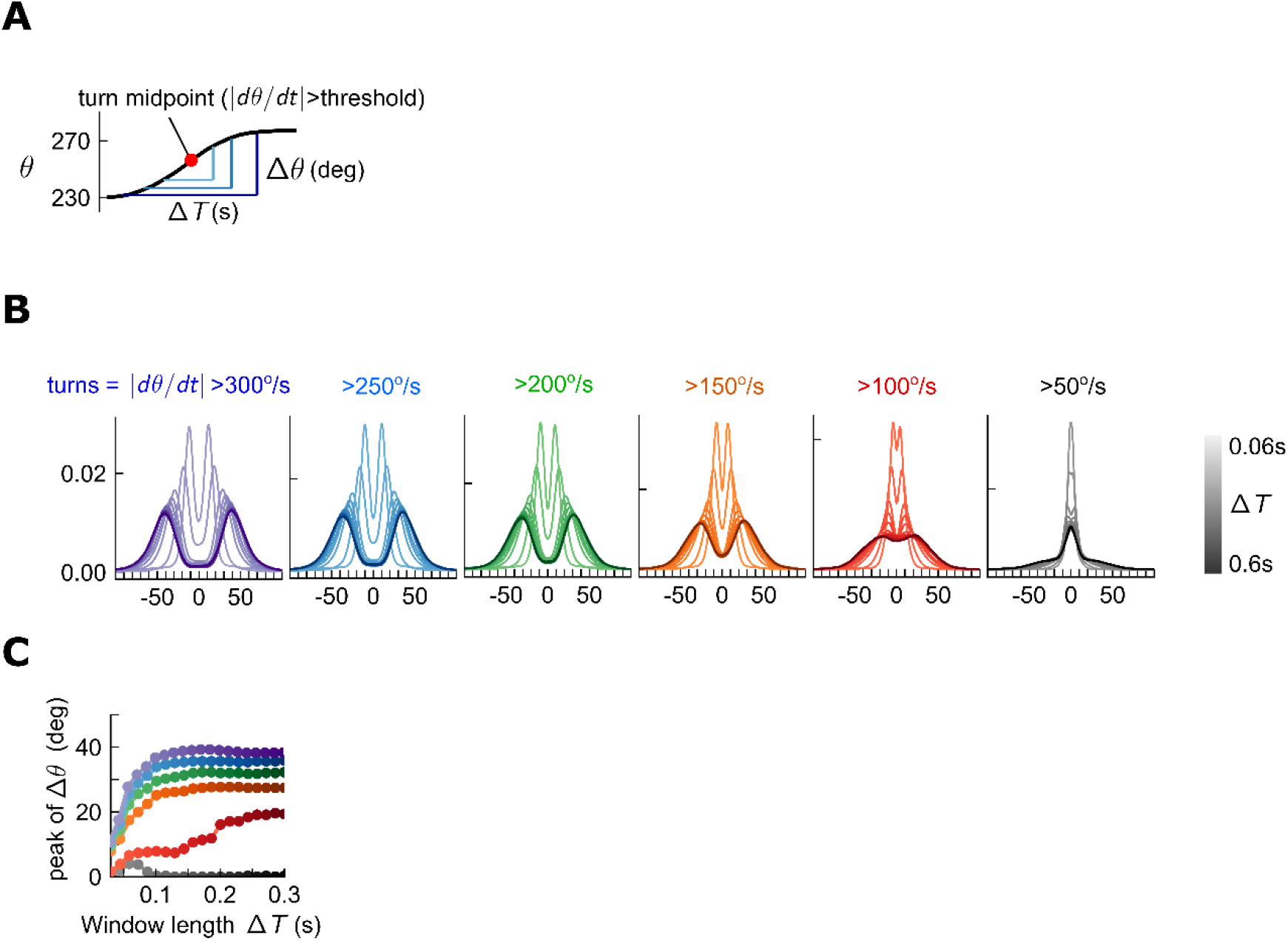
Quantification of turn detection. **A**. Turns occur when angular speed crosses a threshold. The total change in orientation Δ0 is the change in angle over some time window Δ*T* centered on the turn time. **B**. Distributions of orientation changes Δ0 for various time windows Δ*T* (shades) about the turn midpoint, for various thresholds (colors). For all thresholds > 150 deg/s, orientation change Δ0 increases with window Δ*T* at first, but then the distributions remain steady, maintaining a bimodal structure. **C**. Location of peak Δ0 for different turn thresholds, as a function of window length Δ*T*. For all thresholds > 150 deg/s, peak Δ0 remains constant for sufficient Δ*T*.

**Figure S6:**
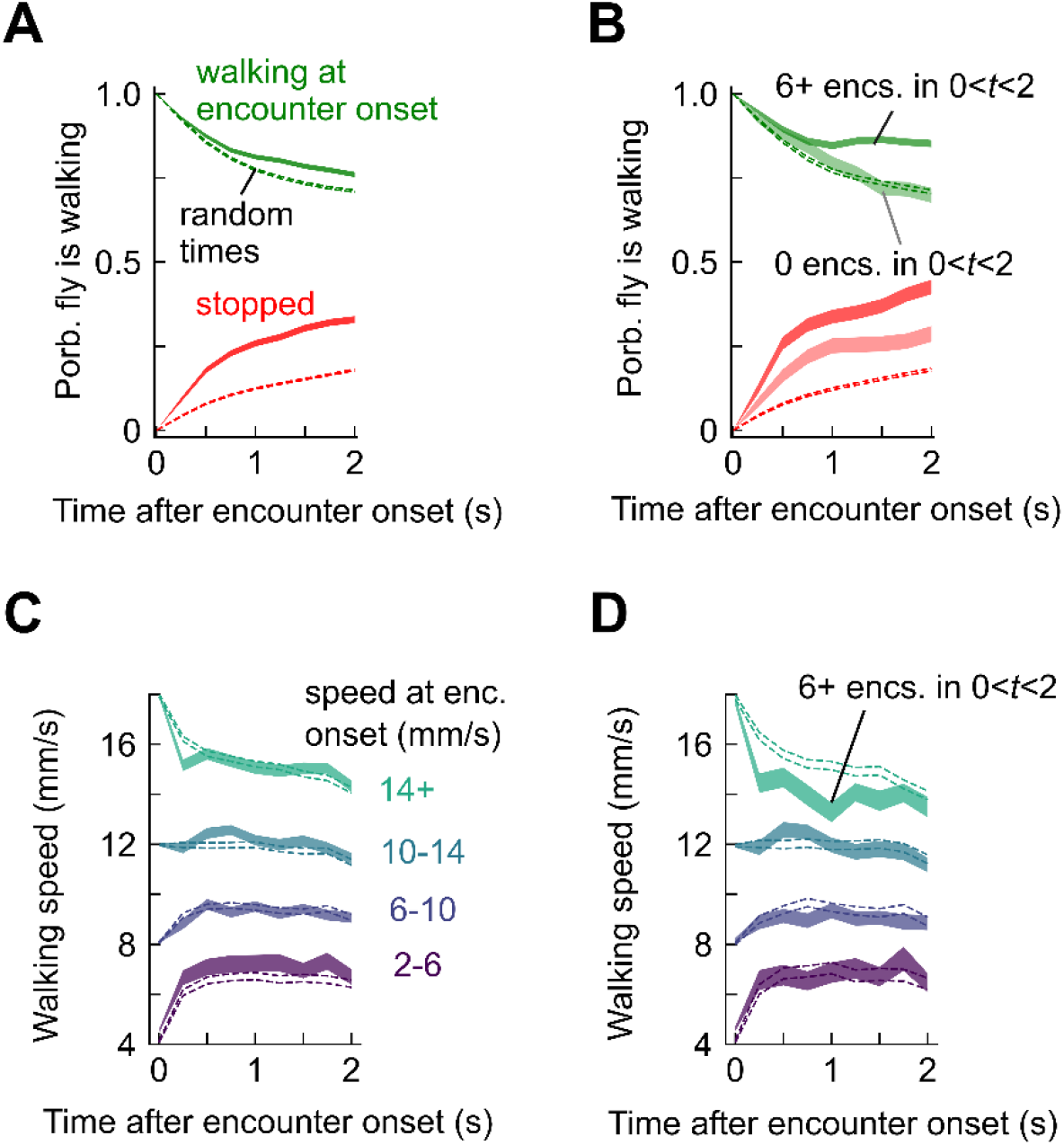
Stop and walk decisions depend on encounter timing. **A. P**robability of walking (green) and being stopped (red) in a 2 second window following an odor encounter (solid) or at random times (dashed). **B**. Same, for encounters with zero subsequent encounters (lighter shade) or >5 subsequent encounters (darker shade) in the following 2 seconds. **C**. Walking speed in a 2s window following encounters, binned by speed at encounter onset. Dotted lines indicate randomly chosen times. **D**. Same, for 6 or more subsequent encounters in the 2s window.

**Figure S7:**
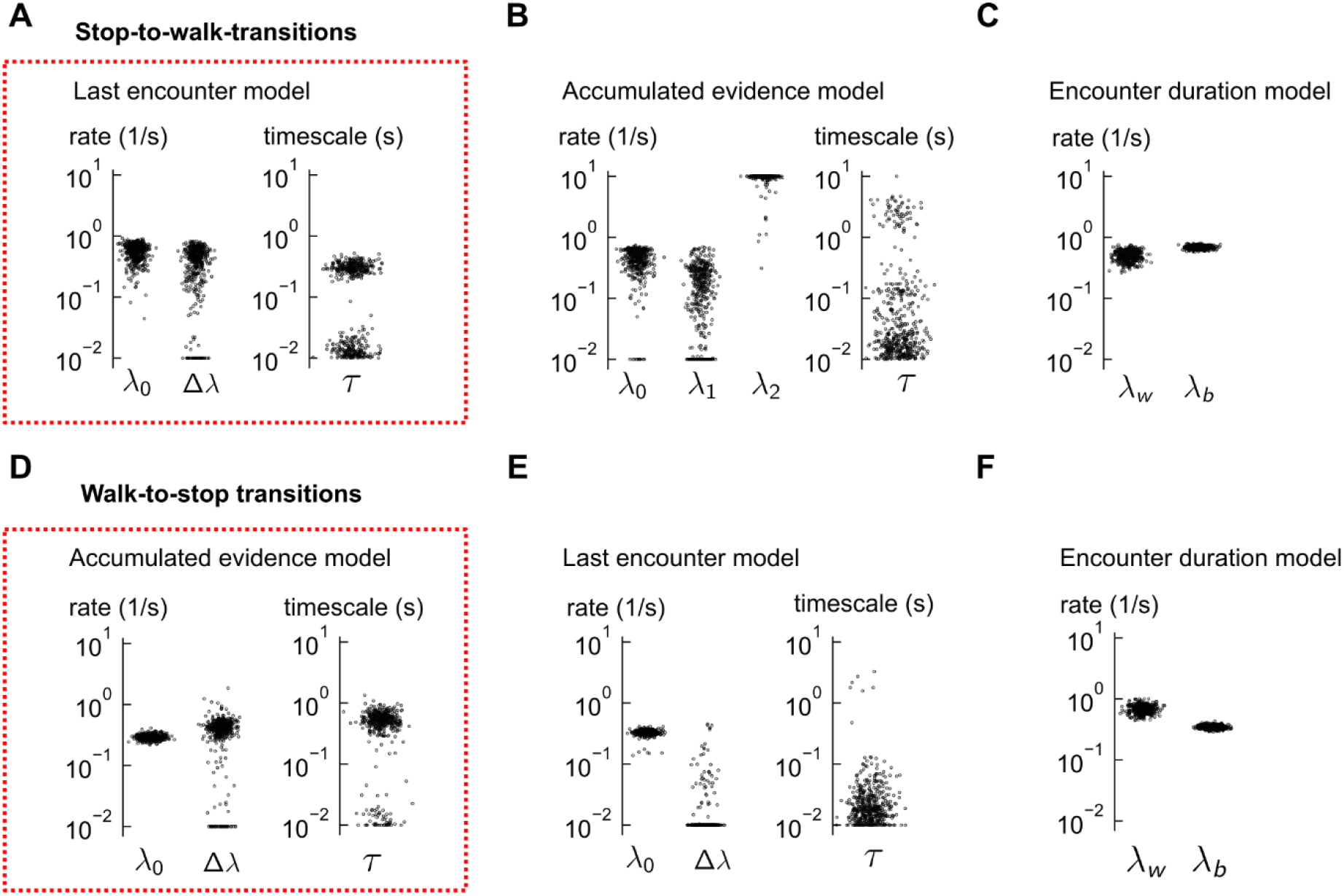
Distributions of estimated parameters for all models. **A.** The estimated parameters for the last encounter model of walk-to-stop transitions, using 500 distinct subsets of the data. Red box indicate this model was the best fit to the data. Occasionally, the global minimum of the nonconvex cost function cannot be found and the optimization terminates near the parameter bounds, rendering the parameter distributions bimodal. **B-C.** Same for the other two models which did not explain the data. In some cases, the global minimum consistently lies at the boundary (e.g. *λ*_2_ in the accumulated evidence model), suggesting that the model is poor. In both models, the predictions were poor (Figure 5H-I). **D**. The estimated parameters for the accumulated evidence model of stop-to-walk transitions, using 500 distinct subsets of the data. **E-F.** Same for the other two models which did not explain the data well. In the last encounter model, the Δ*λ* parameter was consistently estimated at its bound (effectively zero). In both models, the predictions were poor (Figure 6H-I).

**Supplemental Video S1.**

Starved wild-type (CS) female flies navigating in intermittent smoke plume.

**Supplemental Video S2.**

Simultaneous quantification of odor stimulus and fly behavior. Starved wild-type (CS) female flies navigating in intermittent smoke plume. Trajectory of a single fly (same as in Figure 1A) is displayed with its speed, orientation and perceived signal.

**Supplemental Video S3.**

Starved wild-type (CS) female flies navigating in intermittent smoke plume. Smoke valve is turned on/off at 15s blocks while wind is kept fluctuating. Green and blue dashed lines in the orientation panel represent up-wind and cross-wind orientations, respectively.

